# Deficits in mitochondrial function and glucose metabolism seen in sporadic and familial Alzheimer’s disease derived Astrocytes are ameliorated by increasing hexokinase 1 expression

**DOI:** 10.1101/2023.03.23.534020

**Authors:** Simon M Bell, Hollie Wareing, Alexander Hamshaw, Suman De, Elizabeth New, Pamela J Shaw, Matteo De Marco, Annalena Venneri, Daniel J Blackburn, Laura Ferraiuolo, Heather Mortiboys

## Abstract

**Background:** Astrocytes have multiple roles including providing neurons with metabolic substrates and maintaining neurotransmitter synaptic homeostasis. Astrocyte glucose metabolism plays a key role in learning and memory with astrocytic glycogen a key substrate supporting memory encoding. The neuronal support provided by astrocytes has a high metabolic demand. Deficits in astrocytic mitochondrial metabolic functioning and glycolysis could impair neuronal function. Changes to cellular metabolism are seen early in Alzheimer’s disease (AD). Understanding cellular metabolism changes in AD astrocytes could be exploited as a new biomarker or synergistic therapeutic agent when combined with anti-amyloid treatments in AD.

**Methods:** In this project, we characterised mitochondrial and glycolytic function in astrocytes derived from patients with sporadic (n=6) and familial (PSEN1, n=3) forms of AD. Astrocytes were derived using direct reprogramming methods. Astrocyte metabolic outputs: ATP, and extracellular lactate levels were measured using luminescent and fluorescent protocols. Mitochondrial respiration and glycolytic function were measured using a Seahorse XF Analyzer. Hexokinase deficits identified where corrected by transfecting astrocytes with an adenovirus viral vector containing the hexokinase 1 gene.

**Results:** There was a reduction of total cellular ATP of 20% (p=0.05 in sAD astrocytes) and of 48% (p<0.01) in fAD. A 44% reduction (p<0.05), and 80% reduction in mitochondrial spare capacity was seen in sAD and fAD astrocytes respectively. Reactive oxygen species (ROS) were increased in both AD astrocyte types (p=0.05). Mitochondrial complex I and II was significantly increased in sAD (p<0.05) but not in fAD. Astrocyte glycolytic reserve and extracellular lactate was significantly reduced when compared to controls in both sAD and fAD (p<0.05). We identified a deficit in the glycolytic pathway enzyme hexokinase, and correcting this deficit restored most of the metabolic phenotype in sAD but not fAD astrocytes.

**Conclusion:** AD astrocytes have abnormalities in functional capacity of mitochondria and the process of glycolysis. These functional deficits can be improved by correcting hexokinase expression deficits with adenoviral vectors. This suggests that hexokinase 1 deficiency could potentially be exploited as a new therapeutic target for AD.

## Background

Alzheimer’s disease (AD) is the most common cause of dementia worldwide [1]. It is estimated that over 44 million people have the condition across the globe, with numbers of patients with the condition expected to triple by 2050 [2]. The sporadic form of AD (sAD) accounts for the majority of cases of the disease, with familial disease (caused by mutations in the Amyloid precursor protein, Presenilin 1 &2 genes) accounting for approximately 5% of cases [3]. All familial forms of AD (fAD) lead to an increase in the production and deposition of amyloid beta (Aβ) within the brain [4–9]. As amyloid deposition within the brain is also seen in sporadic forms of AD, this led to the development of the amyloid cascade hypothesis for the initiation of the pathogenesis of AD [10–13].

Amyloid and tau aggregates within the brain are clearly an important part of the pathology of AD. However, therapeutics designed to remove amyloid from the brain have shown mixed improvements in clinical outcomes for people with AD [14–16]. The recently licensed Aducanumab, an amyloid clearance monoclonal antibody, showed some success in improving cognitive performance [17]. Lecanemab, another amyloid clearing monoclonal antibody, reduces cognitive decline by 27% in patients with AD and was the first amyloid monoclonal antibody to reach all primary endpoints in clinical trials [18]. As clearing amyloid from the brain alone does not completely stop or reverse the disease course, investigating other pathophysiological processes is important to fully understand the pathological and clinical phenotype of AD [16].

Metabolic changes are seen early within the brain of people with AD, and areas of high glucose metabolism are the same areas affected by Aβ aggregates, tau accumulation, and cortical atrophy [19, 20]. AD pathology prone brain regions show a strong negative correlation between brain metabolism and AD-related gene expression [21]. This has led to the suggestion that metabolic failure, or reductions in metabolic efficiencies of brain cells may be a key step in the development of AD [22].

We and other groups have shown that the nervous system is not the only site of both mitochondrial dysfunction, and glycolytic change in AD, with fibroblasts [23–28], platelets[29] and white blood cells [30] all showing metabolic abnormalities. We have also shown that the parameters of mitochondrial function; mitochondrial spare respiratory capacity (MRSC) and mitochondrial membrane potential (MMP) in fibroblasts correlate with neuropsychological changes seen in AD [25]. This suggests that both central and peripheral metabolism changes have an important role in AD.

Our previous work, and that of others has shown the capacity for both OxPHOS and glycolysis is impaired in the fibroblasts taken from patients with sAD and fAD [24, 25, 28, 31–37]. The changes reported are associated with a reduction in mitochondrial membrane potential (MMP), which may lead to ATP reduction in times of stress. If the metabolic capacity of CNS cells becomes impaired, then this could lead to the development of metabolic failure of the brain in times of increased energy expenditure. Imaging studies undertaken in people with AD suggest that metabolic failure occurs early in AD, as the brain is less able to uptake glucose as the disease progresses [38–42].

Interestingly, several established pathologies interact with the metabolism of glucose in AD. Hexokinase 1, the predominant isoform of the hexokinase enzyme within the brain, is the first enzyme in the glycolytic pathway and is associated with the mitochondrial outer membrane. There is a dramatic effect on hexokinase function when the enzyme stops being bound to the mitochondrial membrane [43, 44]. The amyloid-β peptide has been shown to redistribute hexokinase away from the mitochondria, reducing its activity [45]. Activation of the Glycogen synthase kinase-3 beta enzyme, which causes tau hyperphosphorylation and amyloid accumulation, has also been shown to dislodge hexokinase from the mitochondrial membrane [46, 47].

Interleukin-1β a pro-inflammatory cytokine raised in AD has been shown to reduce the expression of hexokinase and also cause dissociation from the mitochondrial membrane [48]. Therefore metabolism, and specifically the function of the hexokinase 1 enzyme, interacts with the key pathological substrates of AD of protein accumulation and neuroinflammation.

The vast majority of net brain metabolism occurs in neurons and astrocytes. These two cell types form a metabolic relationship, which sees the astrocyte play a pivotal role in maintaining many cellular functions within the neuron [49–52]. Astrocytes provide neurons with metabolic substrates such as lactate, maintain the concentration of ion gradients and neurotransmitters at neuronal synapses, and can divert blood flow to areas with high metabolic activity, allowing increased oxygen and glucose uptake [53–55]. Astrocyte glucose metabolism has a key role in learning and memory with astrocytic glycogen shown to be a key metabolic substrate of memory encoding [56–58]. This dependent relationship between the neuron and the astrocyte puts specific metabolic demands on this cell type, meaning that any abnormalities in the function of astrocyte mitochondria, or the astrocytes ability to metabolise glucose will affect both astrocyte and neuron function.

Multiple studies have shown that pathological changes occur in astrocytes in AD. Altered insulin signalling has been identified [59], as have changes in calcium homeostasis [60, 61], and the ability of astrocytes to metabolise amyloid [62]. Glycolysis-derived metabolites from astrocytes have also been shown to modulate synaptic function [63]. As astrocytes have a key role in neuronal support, memory acquisition, and the AD pathological process, a thorough understanding of their metabolic phenotype may highlight new AD therapeutic targets or biomarkers. Metabolic change in astrocytes has been difficult to study in human AD astrocytes until recently due the lack of living human metabolically active astrocytes available for experimentation.

Over the last 10 years induced pluripotent stem cell (iPSC) technology has revolutionized neuroscience research, as it allows the creation of multiple nervous system cell linages from patients who have the condition under investigation [64–67]. This technology reprograms terminally differentiated human cells, usually fibroblasts, into stem cells, which can then be differentiated into many different cell types. An initial criticism of this technique when applied to neuroscience research was that neuronal lineage cells created shared gene expression profiles, and functions more similar to foetal cells than adult ones [68, 69]. Recent advances in iPSC technology have meant that direct reprogramming of terminally differentiated cells can now be performed without returning cells to a true stem cell phase [70, 71]. This reprogramming method allows cells to maintain some of the aged phenotype of their parent cell. This is an advantage when studying AD, as increasing age is the largest risk factor for developing the disease.

In this paper we use induced neuronal progenitor cell technology [72–74] to derive astrocytes from patients with sAD and fAD (Presenilin 1 mutations). We investigated glycolysis and mitochondrial function, and how this correlates with neuropsychological changes seen early in AD. Finally, we have investigated the role that hexokinase 1 deficiency plays in the development of both mitochondrial functional and glycolysis deficits in AD, and if correcting this deficit improves astrocyte metabolic output.

## Methods

### Patient Details

Sporadic AD (sAD) patients and matched controls were recruited as part of the MODEL-AD study (Yorkshire and Humber Research and Ethics Committee number: 16/YH/0155) for fibroblast biopsy. All sAD and matched controls had previously participated in the European EU-funded Framework Programme 7 Virtual Physiological Human: Dementia Research Enabled by IT (VPH-DARE@IT) initiative (http://www.vph-dare.eu/). A diagnosis of Alzheimer’s disease was made in sAD patients based on clinical guidelines [75]. Table 1 shows the demographic details of the sAD patients.

**Table 1.** Patient Demographic Information and contemporary treatment status. #For control 7 some items of the MMSE could not be tested due to sensory impairment.

| Patient Line | Age (Years) | Sex | MMSE | Length of Education (Years) | AD Treatment (at biopsy) |
| --- | --- | --- | --- | --- | --- |
| 1 | 60 | Male | 18 | 9 | None |
| 2 | 59 | Female | 23 | 11 | Galantamine |
| 3 | 63 | Female | 26 | 10 | Donepezil |
| 4 | 60 | Male | 18 | 11 | Memantine |
| 5 | 60 | Male | 18 | 11 | None |
| 6 | 79 | Female | 28 | 15 | None |
| Group Mean (Standard Dev) | 63.5 (7.71) |  | 21.83 (4.48) | 11.16 (2.04) |  |
| Control lines |  |  |  |  |  |
| 1 | 52 | Male | 27 | 16 | NA |
| 2 | 61 | Female | 30 | NA | NA |
| 3 | 100 | Female | 24# | 14 | NA |
| 4 | 52 | Female | 29 | 17 | NA |
| 5 | 56 | Male | 24 | 11 | NA |
| 6 | 73 | Female | 26 | 12 | NA |
| 7 | 75 | Female | 28 | 18 | NA |
| Group Mean (Standard Dev) | 66.7 (17.3) |  | 27.33 (2.16) | 14.66 (2.80) |  |

Presenilin 1 AD patient fibroblasts (fAD) and matched controls were acquired from the NIGMS Human Genetic Cell Repository at the Coriell Institute for Medical Research: (ND41001, ND34733, AGO6848, GMO2189), and a cohort from a Sheffield based study (155 and 161). All fAD patients had a confirmed pathological mutation in the presenilin 1 gene (See Table 2)

**Table 2.** Patient demographics for Familial AD presenilin 1 lines and associated controls.

| Control Line | Age<br>(Years) | Sex | Presenilin 1 Mutation |
| --- | --- | --- | --- |
| Sheffield 161 | 31 | Male | NA |
| Sheffield 155 | 40 | Male | NA |
| Coriell (GM02189) | 63 | Male | NA |
| Group Mean<br>(Standard Dev) | 44.66<br>(16.5) |  |  |
| Familial Line | Age<br>(Years) | Sex | Presenilin 1 Mutation |
| Coriell (ND41001) | 47 | Female | 14q24.3<br>Intron 4, G deletion |
| Coriell (ND34733) | 60 | Male | P264L |
| Coriell (AG06848) | 56 | Female | missense mutation {Ala246Glu (A246E)} |
| Group Mean<br>(Standard Dev) | 48.33<br>(11.06) |  |  |

### Tissue Culture

Fibroblasts were cultured in a DMEM based media (Corning) with sodium pyruvate (1%) [Sigma Aldrich], penicillin & streptomycin (1%) [Sigma Aldrich], and Fetal Bovine Serum (10%) [Biosera], incubated at 37^0^C in a 5% carbon dioxide atmosphere.

### Fibroblast Reprogramming

Fibroblast reprogramming into Induced Neuronal Progenitor Cells (iNPCs) and astrocyte differentiation was carried out as previously described, [71]. In brief, fibroblasts were plated at a density of 250,000 cells per well. 24 hours after plating fibroblasts were transduced with lentiviral non-integrating vectors (OCT3, Sox2, and KLF4,). 48 hours after transduction, cells were changed into iNPC media consisting of DMEM/F12, 1% N2, 1% B27, EGF (40 ng/ml) and FGF (20 ng/ml). Once iNPCs lines were established expression of the neuronal progenitor markers PAX-6 and Nestin was confirmed through immunohistochemistry. iNPCs were maintained for ∼30 passages.

Astrocytes were differentiated from iNPCs using established protocols (ref). Briefly, iNPCs were seeded on a 10cm dish coated with fibronectin (5 µg/ml, Millipore) in DMEM media (Lonza) containing 10% FBS (Biosera), and 0.3% N2 (Gibco) and differentiated for 7 days. After differentiation expression of several astrocyte cell markers was confirmed using immunohistochemistry (details described in section below). This included GFAP, Vimentin, CD44, ALDHL1 and NDGR2. A key functional ability of astrocytes is the ability to uptake glutamate. This was confirmed in our astrocyte cell populations by performing a glutamate uptake assay (see below). We have also previously published evidence highlighting that astrocytes differentiated in this way maintain an aged cell phenotype [76].

### Immunohistochemistry

Astrocytes or iNPCs were plated on a black 96-well plate (Greiner bio-one) at a density of 2500 cells/well. After 24 hours cells were fixed with 4% paraformaldehyde (PFA) for 10 minutes. Cells were permeabilised in a solution of 0.1% triton, PBS and TWEEN 20 (1:1000) (PBST) for 30 minutes. Following this, cells were washed with PBST twice. Cells were then blocked in PBST and horse serum (5%) for 60 minutes. Fixed cells were incubated with primary antibodies overnight (Supplementary table 1). Secondary antibodies were added at a concentration of 1:1000 for 1 hour, then Hoechst dye (Life Technologies) at a concentration of 2.5µM for 10minutes. Staining was visualised using an OPERA high content imager (Perkin Elmer).

### Glutamate Uptake Assay

An Abcam colorimetric Glutamate Assay kit (ab83389) was used to measure astrocyte glutamate uptake. The assay was performed as per manufacturer’s instructions. In brief, astrocytes were plated on a black 96 well plate at a density of 10,000 cells/well at day 5 of differentiation. On day 7 of differentiation the astrocyte media was changed to Hank’s Balanced Salt Solution (HBSS) (Gibco), without calcium or magnesium for 30 minutes. This media was then changed to HBSS containing magnesium and calcium for 3 hours which also contained 100µl of glutamate at a concentration of 1:1000. Samples were then collected as described by manufacturer and snap frozen in liquid nitrogen. Glutamate measurement was performed from this point onwards as per kit protocol. Percentage glutamate uptake was then calculated based on the known concentration of glutamate added to the astrocytes.

### Total Cellular ATP levels & ATP Inhibitor Assay

Cellular Adenosine Triphosphate (ATP) levels were measured with the ATPlite kit (Perkin Elmer) as previously described (ref). Astrocytes were plated at a density of 5000 cells per well in a white Greiner 96 well plate on day 5 of differentiation. ATP levels were corrected for cell number using CyQuant (ThermoFisher) kit, as previously described [25], and then disease lines were normalized to controls. The same assay was performed using the inhibitors of glycolysis (2-deoxyglucose 50mM, [Sigma]) and OxPHOS (Oligomycin 1µM, [Sigma]) to assess the reliance on each metabolic pathway for total cellular ATP. A glycolysis and OxPHOS inhibitor assays were performed. Oligomycin (OxPHOS inhibitor) or 2-deoxyglucose (glycolysis inhibitor), or both inhibitors were added to the cells for 30 minutes and incubated at 37^0^C during this time. The ATP assay was then performed as described above.

### Mitochondrial Membrane Potential

Astrocytes were plated at a density of 2500 cells per well in a black Greiner 96-well plate at day five after the start of differentiation. On day 7 of differentiation cells were incubated with tetramethlyrhodamine (TMRM) for 1 hour (concentration 80nM) and Hoechst (concentration 10nM) (Sigma Aldrich). Cells were incubated at 37^0^C during this time. Dyes were then removed, and astrocytes were maintained in MEM media whilst imaging using an InCell Analyzer 2000 high-content imager (GE Healthcare). 25 fields with an average number of 500 cells per well at an emission/excitation spectrum of 543/604m was imaged. Mitochondrial morphological parameters and cell area were quantified using an INCELL developer protocol[77]. Parameters assessed include mitochondrial form factor. This is a combined measurement of both a mitochondria’s perimeter length and area (Form Factor=1/(Perimeter^2^/4π.area). It is a measure of how round a mitochondrion appears to be, but also gives an indication of how interconnected the mitochondrial network is.

### Extracellular Lactate Measurement

Extracellular lactate was measured using an L-Lactate assay kit (Abcam ab65331). At day 7 of differentiation 1µl of media was removed from a 10cm dish containing confluent astrocytes and used in the assay as per the manufacturer’s instructions. Briefly, 1µl of media was added to 49µl of lactate assay buffer, to this 50µl reaction mix was then added and the samples are then incubated at room temperature for 30minutes. Lactate measurements are then made on a PHERAstar Plate reader absorbance filter at 570nm. To calculate the lactate concentration a standard curve was measured (2-10nmol range of lactate concentrations), and background lactate measurements were made. Measurements were normalized to controls on each separate day of experimentation.

### ATP Substrate Assay

The ATP substrate assay was used to investigate mitochondrial ATP production in the presence of complex I and II substrates. 500,000 where cells are collected at day 7 of astrocyte differentiation. Methods have been previously described by Manfredi et al 2002, [78]. In brief, cells are suspended in 250µl of buffer A (KCL 150mM, Tris HCL 25mM, EDTA 2mM, BSA 0.1%, K_3_PO_4_ 10mM, and MgCL 0.1mM, pH 7.4). Cells were then permeabilised with histone 2ug/ml for 5 minutes. After permeabilization 5 volumes of buffer A (1250mls) was added to the cell suspension. The suspension was then centrifuged for 5 minutes at 17,000g. Cells were then resuspended in 150µl of Buffer A. 550µl of buffer A was added to the remaining 100µl of suspension for use in the substrate assay. The assessment of total cellular ATP at this stage must be performed with 15minutes of adding the 550µl of buffer A to samples.

A PHERAstar plate reader (BMG Labtech) in luminescence mode was used. A background luminescence reading is made of each well on the assay plate containing 160µl of cell suspension then one of either the complex I substrates (malate 1.25mM and galactose 1.25mM), or complex II substrates (succinate 1.25mM, rotenone 1µM added as complex I inhibitor) is added. After baseline kinetics were measured the machine was paused and 5µls of Adenosine diphosphate (ADP, 4µM) and 10µl of the ATP substrate solution, described above in the ATP assay section were added to each well. The kinetics assay is then resumed, and measurements of substrate use are made for approximately the next 30 minutes. The gradient of the kinetics curve is then calculated and normalised to protein content using a Bradford assay. Disease samples are normalised to controls of the day.

### Metabolic flux assay

#### Mitochondrial Stress Test

Astrocyte OxPHOS was assessed by measuring oxygen consumption rates (OCR) using a 24-well Agilent Seahorse XF analyzer. Astrocytes were plated at a density of 10,000 cells per well at day 5 of differentiation. At day 6 of differentiation cells were either switched to a galactose containing media or continued to be maintained in glucose. At day 7 of differentiation astrocytes were switched to XF media (Agilent) and then assessed using the previously described *Mitochondrial Stress Test Protocol* [24]. During this protocol measurements were taken at the basal point: after the addition of oligomycin (0.5µM), after the addition of carbonyl cyanide-4-(trifluoromethoxy) phenylhydrazone (FCCP) (0.5µM) and after the addition of rotenone (1µM). OCR measurements were normalized to cell count, as previously described [24]. ATP linked respiration, Mitochondrial respiration, Mitochondrial Spare Respiratory Capacity, Respiratory control ratio and ATP couple respiration are all calculated from this experiment.

#### Glycolysis Stress Test

Astrocyte glycolysis was measured using the glycolysis stress test protocol on a 24-well Agilent Seahorse XF analyzer. Astrocytes were plated at a density of 10,000 cell per well at day 5 of differentiation. At day 7 of differentiation as with the mitochondrial stress test astrocytes were transferred to XFmedia (Agilent) and glycolysis was assessed. The glycolysis stress test was used to assess glycolysis, this has been described previously, [25]. In brief, three measurements were taken at the basal point: after the addition of glucose (10µM), after the addition of oligomycin (1.0µM) and after the addition of 2-deoxyglucose (50µM). Basal rate of glycolysis, Glycolytic capacity, Glycolytic reserve and non-glycolytic acidosis are all measured during the glycolysis stress test. Measurements were normalized to cell count, as previously described [24]

### Glucose Uptake

Glucose uptake of astrocytes was measured using the Glucose Uptake Assay Kit (Fluorometric) (Abcam, ab136956). Astrocytes were plated at a density of 2500 cells per well in a 96 well black Greiner plate on day 5 of differentiation. On day 7 of differentiation the astrocytes were incubated for 40 minutes in a 100µl of Krebs-Ringer-Phosphate-Hepes (KRPH) buffer with 2% bovine serum albumen added pH 7.4. Astrocytes were then treated with 2-deoxyglucose at a concentration of 50mM for 20minutes suspended in KRPH (volume 100µl per well). Cells were then treated with 90µl of Extraction buffer (Abcam), snap frozen in liquid nitrogen, and then thawed on ice. 25µl of sample was then placed in a black 96-well plate, with the volume made up to 50µl with glucose uptake assay buffer (Abcam). The assay from this point onwards was performed according to the manufacturer’s instructions. Briefly, using the 2-deoxyglucose standard a standard curve was created with concentrations between 0µM-20µM of 2-Deoxyglucose. 50µl of each standard was plated on the black 96-well plate. 50µl of reaction mix (47µl Glucose uptake assay buffer, 2µl of enzyme mix and 1µl of PicoProbe, all supplied by Abcam) was added to each sample and standard well. Samples were then incubated for 40 minutes at 37^0^C. Measurements of Fluorescence at Ex/Em=535/587nm were then taken using PHERAstar plate reader (BMG Labtech).

### Glutamine/Glutamate assessment

A Glutamine/Glutamate-Glo™ Assay (Promega) was used to measure intracellular concentrations of glutamine and glutamate. A 10cm dish of Astrocytes (≈ 2,000,000 cells) was immersed in inactivation solution (2mls of HCL 0.3N and 1ml PBS) for 5 minutes on day 7 of differentiation. After this the dish was scraped and 1ml of Tris solution (2-Amino-2-(hydroxymethyl)-1,3-propanediol, 450mM at pH 8.0) was added to the cells. 200µl of this solution was then added to 200µls of PBS. 25µl of this dilution is then placed in a well of a white 96-well Greiner plate. After sample preparation the assay was performed as per the protocol provided by Promega.

### Mitochondrial Complex Assays

Mitochondrial complex assays were performed for mitochondrial complexes I (ab109721), II (ab109908) and IV (ab109909). Assay kits were obtained from Abcam, and the standardised protocol offered with the kit was followed. Approximately 2,000,000 astrocytes per experiment were used for each complex assay round. Once complex assay values had been assessed values where normalised to sample protein content using a Bradford assay and then the controls assessed on the day of the experiment.

### Hexokinase Activity Assay

Hexokinase activity was measured using the Abcam (ab136957) Hexokinase Assay Kit (Colorimetric). Standardised protocols as described in the kit protocol booklet where followed. Astrocytes were harvested on day 7 of differentiation. In brief, approximately 2,000,00 astrocytes were collected and homogenized using the assay buffer. Samples where centrifuged at 12,000 rpm, for 5 minutes and then kept on ice until assay assessment. Once complex assay values had been assessed values where normalised to sample protein content and then the controls assessed on the day of the experiment.

### Quantitative polymerase chain reaction (qPCR)

For qPCR approximately 2,000,000 astrocytes were harvested. The RNeasy Mini Kit (Qiagen) was used to extract RNA from samples. RNA extraction was performed as described in the kit protocol. Supplementary Table 2 displays the forward and reverse primer sequences used for each gene assessed. Disease samples were normalised to the controls of the day.

### Mitochondrial reactive oxygen species assays

Mitochondrial reactive oxygen species (ROS) were quantified using the mitochondrially-targeted fluorescent redox sensor NpFR2 [79]. Astrocyte cells were plated at a density of 2500 cells per well on a 96 well plate and incubated with the NpFR2 for 30 minutes prior to imaging. Mitochondrial ROS was visualised using an OPERA high content imager (488 nm excitation, 530/30 nm emission) (Perkin Elmer).

### Western Blots

Westerns were performed as previously described [24] 20µg of protein was loaded per sample. Membranes were probed for Glucose transporter 1, 2 and 4, and Hexokinase 1. Membranes were also probed for GAPDH which was used as a loading control. Supplementary Table 1 primary and secondary antibodies used for western blot analysis.

### Viral transduction

On day 4 of astrocyte differentiation; astrocytes were transduced with hexokinase adenoviral vector (AVV). The hexokinase containing AVV was purchased from VectorBiolabs (RefSeq#: BC008730). All hexokinase AAV experiments were performed with a multiplicity of infection (MOI) of 40. Astrocytes were transduced with the virus for 72 hours and then measurements of mitochondrial and glycolytic function were made. An AVV containing a scramble gene was also plated for each experiment. This was acquired from VectorBiolabs with a MOI of 40 also used for these experiments.

### Metabolic neuropsychological correlations

Correlations between metabolic parameters and neuropsychological tests were performed as described previously [25].

### Statistical Analysis

Statistical analysis for this data set is the same as detailed in [25]. In brief metabolic comparisons between AD group and control groups are compared using t-tests. A Pearson correlation was performed for metabolic neuropsychological correlations. Statistics were calculated using the GraphPad v8 software and IBM SPSS statistics version 25.

## Results

### Patient Demographic Details

sAD patients from which astrocytes were derived had a mean age of 63.5 (SD7.71) years (3 Female) sAD controls had a mean age of 66.6 (sd17.3) years (5 female). Table 1 details the Age, MMSE and length of education for both sAD and control astrocytes. For the fAD group the mean age was 48.33 years (1 female). The controls for this group had a mean age of 44.66 years (3 Male). Table 2 details the age, sex, and PSEN1 mutations of PSEN1 and associated control lines.

### iNPC derived sporadic and familial AD astrocytes display astrocyte markers and functional properties. Sporadic and familial AD astrocytes have reduced total cellular ATP and extracellular lactate

This is the first study to reprogram sAD and fAD patient fibroblasts to iNPCs and differentiate to astrocytes [71, 72]. As this is the first study to use this particular method of iNPC reprogramming to derive AD astrocytes, we assessed the derived cells for expression of known astrocytic/iNPC markers and functional properties. iNPCs from control, sAD and fAD patients all expressed the neuronal precursor markers Nestin and PAX6 (See figure 1A). Astrocytes were assessed with a combination of astrocytic markers, [74, 80, 81]; all astrocyte lines expressed CD44, GFAP, Vimentin, NDGR2, and ALD1L1 (Figure 1B). Figure 1C highlights the major steps of the reprogramming procedure used in this study. Astrocyte cell area was assessed, showing that both sAD (mean area 1848µm^2^ SD± 355.5, p=0.0316) and fAD astrocytes (mean area 1319µm^2^ SD± 36.04, p=0.0424) had a smaller cell area than control astrocytes (Sporadic controls mean area 2953µm^2^ SD± 263.7, familial controls mean area 2206µm^2^ SD± 299.7) (Figure 1D).

**Figure 1.**
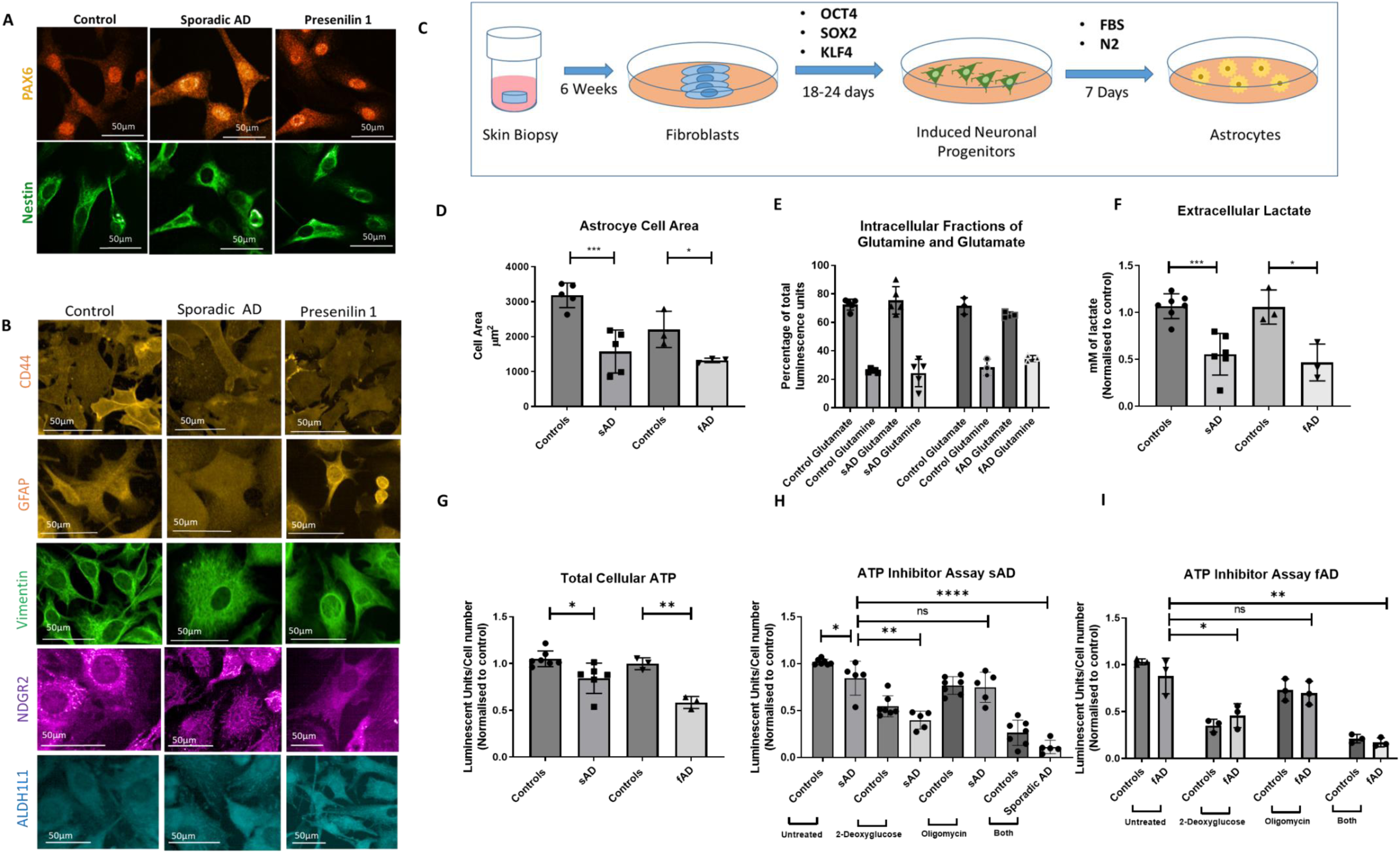
Astrocyte characterization. **A** Control, sporadic and familial AD astrocytes all express the neuronal progenitor cell markers PAX-6 and Nestin, scale bar 50um. **B** Control, sporadic and familial AD astrocytes display staining for CD44 (orange), GFAP (orange), Vimentin (green), NDGR2 (purple), and ALDH1L1 (blue) stained at day 7 of differentiation, scale bar 50um. **C** this panel shows the process of generating the derived astrocytes. Skin biopsies are taken and cultured to generate fibroblasts over approximately 6 weeks. Fibroblasts are then transfected with adenoviral non-integrating vectors (OCT3, Sox2, and KLF4). Induced neuronal progenitor cells are generated over a period of 18-24 days, these cells are then cultured in a medium containing FBS and N2 which leads to the establishment of astrocyte cells after 7 days of differentiation. Panels D-I graph various parameters in astrocytes from sAD and controls, fAD and controls. Each experiment included 7 sAD controls, 6 sAD lines, 3 fAD control lines and 3 fAD lines. Data was analysed after at least 3 technical repeats and across 3 biological repeats were performed in each experiment. Each dot represents the mean of each line across 3 biological repeats. **D** astrocyte cell area. *=p<0.05 ***<u>E</u>*** Intracellular fractions of glutamate and glutamine *=p<0.05. **F** Astrocyte extracellular lactate levels. *=p<0.05 and ***=p<0.001. **G** Total cellular ATP levels *=p<0.05 and **=p<0.01. **H&I** <u>Total cellular ATP determination in the presence of 2-deoxyglucose and</u> <u>oligomycin</u> *=p<0.05, **=p<0.01 and ****=p<0.0001. In all experiments AD astrocytes are compared to relative controls using t-tests.

To assess if the derived astrocytes have key functional astrocytic properties concentrations of intracellular glutamate and glutamine were assessed. Glutamate uptake from the synaptic cleft is a key functional property of in vivo astrocytes [82], and is a key measure in showing *in vitro* astrocytes are functionally active [83, 84]. Control, sAD and fAD astrocytes all had similar intracellular proportions of glutamine and glutamate (Figure 1E). No significant differences were seen between controls and AD astrocytes with regard to intracellular glutamate and glutamine amounts. Extracellular lactate is a key metabolite shared between neurons and astrocytes; therefore we assessed lactate release in the AD astrocytes. Extracellular lactate levels were reduced in both sAD (48% reduction SD 10%, p=0.0003) and fAD astrocytes (59% reduction 15%, p=0.0187) when compared to controls (Figure 1F).

As maintaining energy homeostasis is a key function of astrocytes, we measured total cellular ATP levels. A significant reduction in total cellular ATP was seen in both sAD (19.8% reduction SD6.57%, p=0.0126) and fAD (42% reduction SD 5.2%, p=0.0014) astrocytes when compared to controls (Figure 1G). In order to determine if astrocytes were reliant on glycolysis or OXPHOS for generating ATP, we undertook ATP measurements in the presence of various inhibitors to measure this. A greater proportion of total cellular ATP was produced via glycolysis when compared to OxPHOS in all astrocytes from all groups (Figures 1G). Astrocyte total cellular ATP had a mean reduction of 49% across all three astrocyte groups when glycolysis was inhibited (Control 48%, Sporadic 52% and familial 47% reduction), whereas a mean reduction of 18.4% in total cellular ATP was seen when oligomycin was added to inhibit OXPHOS (Controls 24%, Sporadic 11.4% and familial 20% reduction). These findings reflect the known metabolic dependency of astrocytes on glycolysis for ATP production and provides further characterization of this model (Figures 1H&I).

### AD astrocytes have an altered mitochondrial morphology and MMP

After characterising the AD astrocytes and determining ATP levels, we next assessed mitochondrial morphology and function. sAD astrocytes had a significant reduction in their MMP when compared to controls (22% reduction SD9.9%, p=0.05), whereas fAD astrocytes had a higher MMP then controls (30% increase, p=0.041) (Figure 2A). Mitochondrial morphology was also altered in AD, with both sAD and fAD astrocytes showing an increased number of elongated mitochondria (Sporadic Astrocytes 6.4% increase, p= 0.002, familial astrocytes 16% increase, p=0.0127), and a significantly reduced form factor which due to the equation used to calculate this indicates the mitochondrial network is more interconnected. (Sporadic astrocytes 4.7% SD1% reduction, p= 0.029, familial astrocytes 10% reduction, p=0.036) (Figures 2B&C).

**Figure 2.**
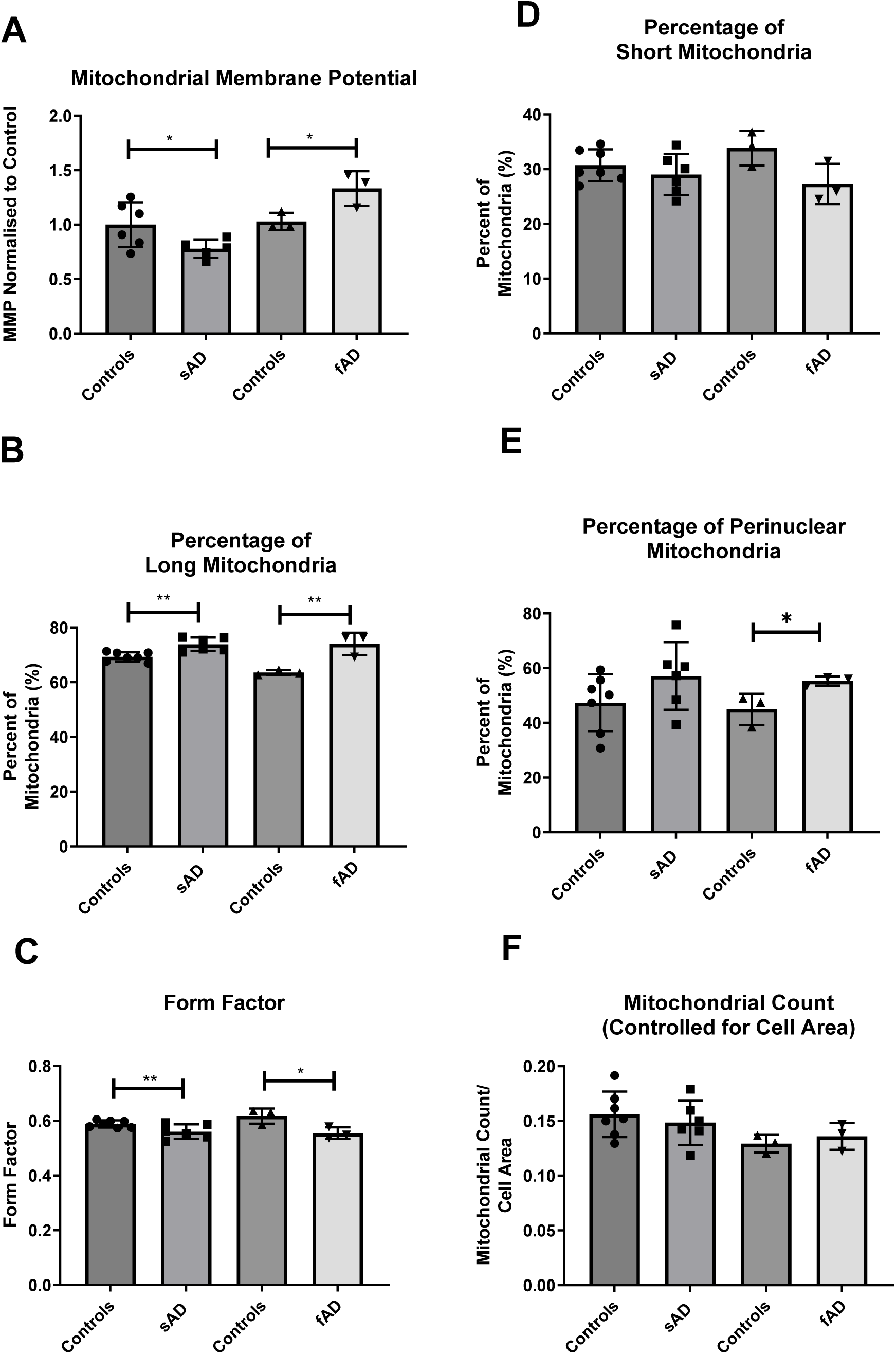
Astrocyte Mitochondrial Function and Morphology. **A** Astrocyte Mitochondrial Membrane Potential. *p<0.05. **B** Percentage of Long Mitochondria as a proportion of the whole mitochondrial. **=p<0.01. **C** Form Factor.*p<0.05 and **=p<0.01.**D** Perinuclear Mitochondria percentage as a proportion of the whole mitochondrial network. *=p<0.05. In all experiments AD astrocytes are compared to relative controls using t-tests. Each experiment included 7 sAD controls, 6 sAD lines, 3 fAD control lines and 3 fAD lines. Data was analysed after at least 3 technical repeats and across 3 biological repeats were performed in each experiment.

A trend towards an increase in the percentage of perinuclear mitochondria was seen in sAD astrocytes (p=0.150), with fAD astrocytes showing a significant increase in perinuclear mitochondrial (23% increase SD 6.93%, p=0.039) when compared to controls (Figure 2D). The percentage of small mitochondria in the mitochondrial network, and mitochondrial number was also measured, but no differences were seen in both Alzheimer types.

### Oxidative Phosphorylation is altered in sporadic and familial AD astrocytes, with a specific reduction in mitochondrial spare respiratory capacity

After showing that AD astrocytes had mitochondrial morphological changes and lower total cellular ATP levels, we next assessed if respiration which is a measure of OxPHOS function was also impaired in AD astrocytes. Figures 3A&B show Oxygen consumption rates (OCR) for sporadic and familial AD astrocytes respectively.

**Figure 3.**
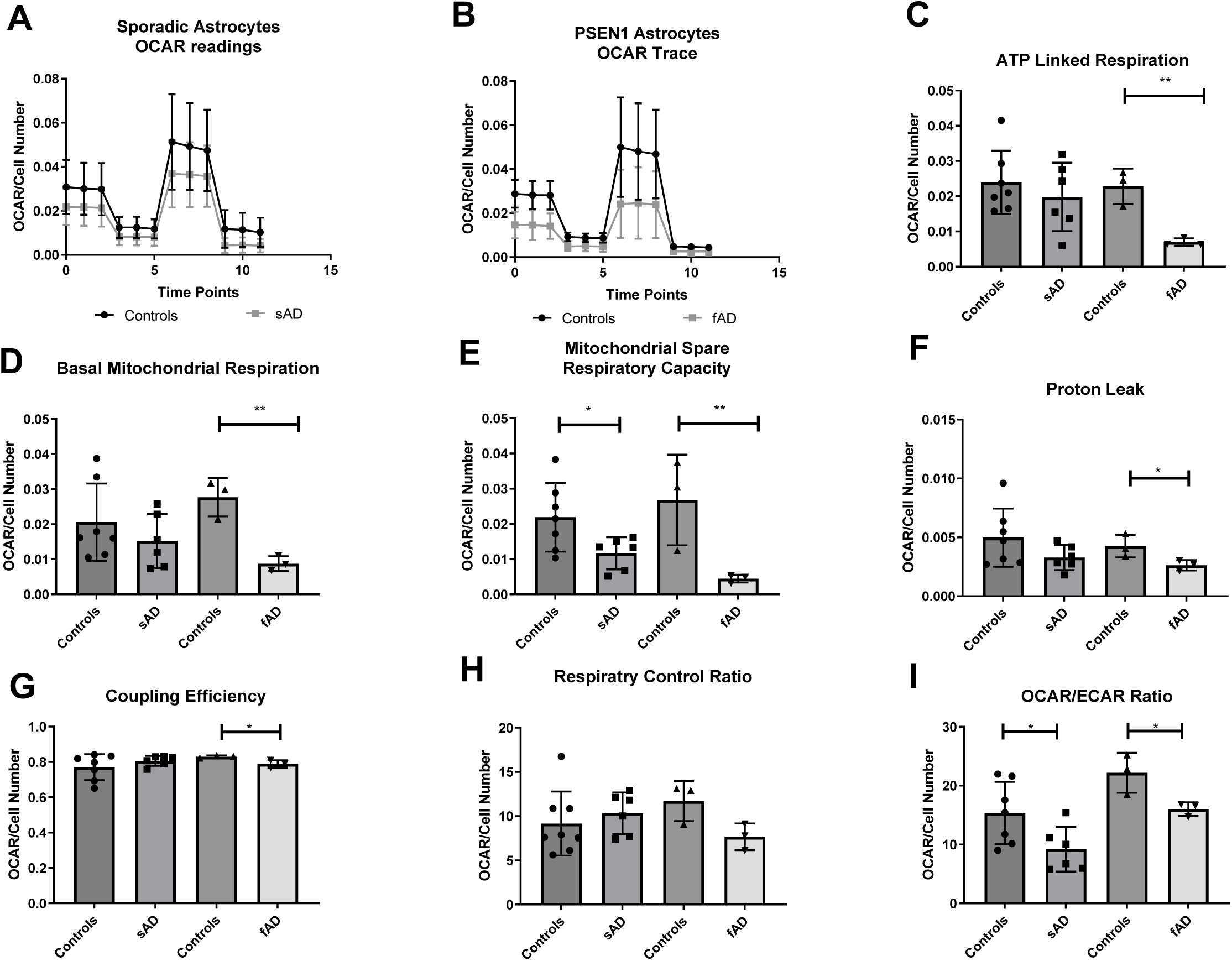
Astrocyte Oxidative Phosphorylation Assessment. **A** Sporadic AD Astrocyte OCR trace, **B** Familial AD astrocyte OCR trace. **C** ATP Linked Respiration. **=p<0.01 **D** Basal Mitochondrial Respiration Basal. **=p<0.01 **E** Mitochondrial Spare Respiratory Capacity. *=p<0.05 and **=p<0.01 **F** Proton leak. *=p<0.05. **G** Coupling Efficiency. *=p<0.05. **H** Respiratory Control ratio **I** OCR/ECAR Ratio *=p<0.05, **=p<0.01. In all experiments AD astrocytes are compared to relative controls using t-tests. Each experiment included 7 sAD controls, 6 sAD lines, 3 fAD control lines and 3 fAD lines. Data was analysed after at least 3 technical repeats and across 3 biological repeats were performed in each experiment.

ATP linked respiration (69% reduction SD13%, p=0.005) and Basal mitochondrial respiration (68.4% reduction SD12%, p=0.004) were both significantly lower in fAD astrocytes when compared to their controls (Figures 3C&D). A trend to a reduction in basal mitochondrial respiration (reduction 23.2%, p=0.3405) and ATP linked respiration (reduction 17.2%, p=0.445) was seen with sAD astrocytes, but this was not significant (Figures C&D). MSRC was shown to be significantly reduced in both sAD(46.6% reduction SD19.8%, p= 0.038) and fAD astrocytes (83.5% reduction SD27.8%, p=0.040) when compared to controls (Figure 4E). fAD astrocytes showed a significant reduction in proton leak when compared to controls, (38.5% reduction SD21.8%, p= 0.050), whereas sAD astrocytes only showed a trend towards a reduction (Figure 4F). No significant difference was seen in coupling efficiency or respiratory control ratio in sporadic AD astrocytes when compared to controls (Figures 4G&H), but a significant reduction in coupling efficiency was seen in fAD astrocytes (5% reduction SD 1.4%, p= 0.032, see Figure 4G). These data together suggest that fAD astrocytes have a greater compromise of OxPHOS than sAD astrocytes, although both sporadic and familial AD astrocytes have a significant reduction in MSRC suggesting mitochondrial functional impairment when under conditions of stress.

**Figure 4.**
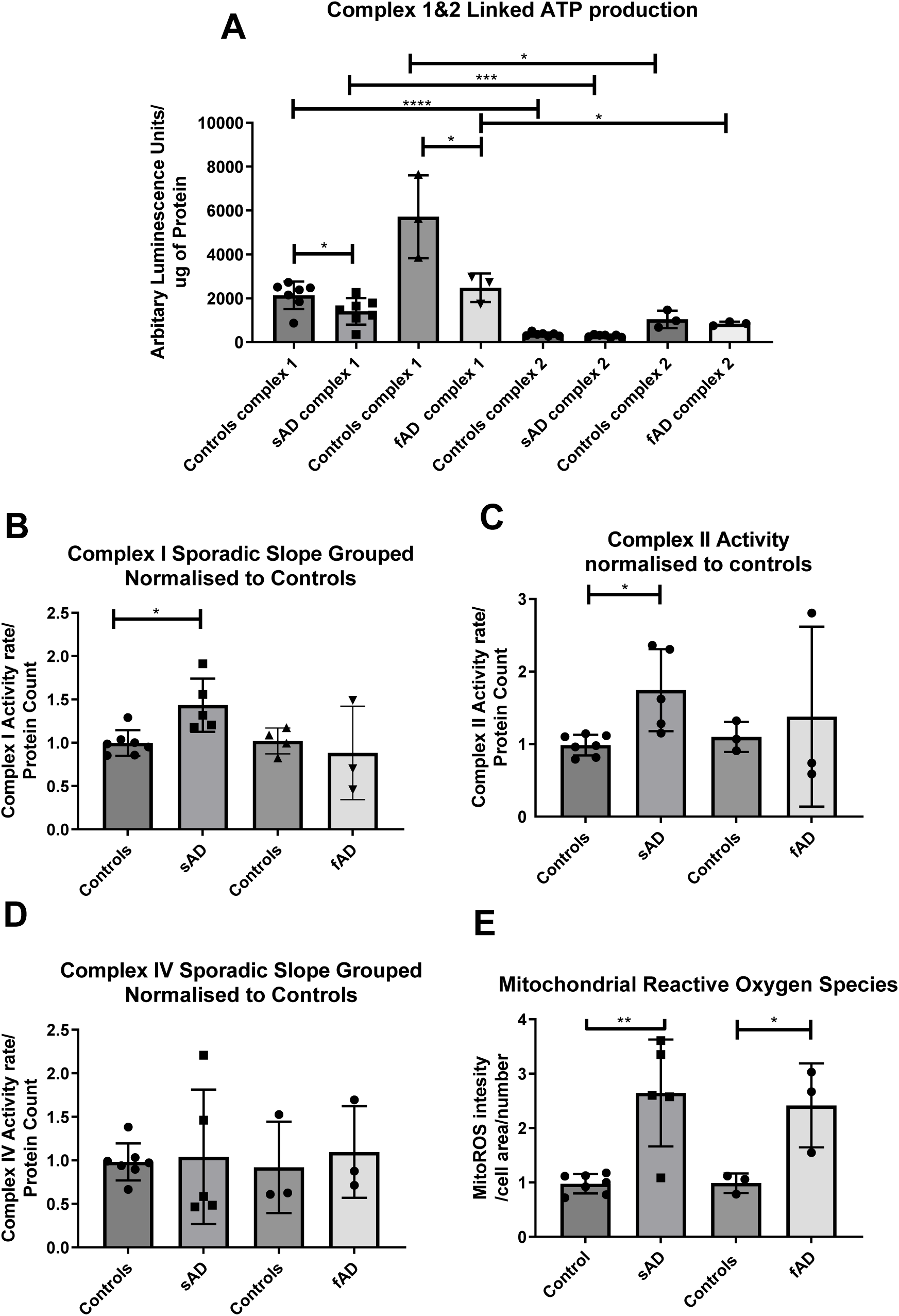
Mitochondrial Electron Transport Chain complex assessment. **A** Assessment of mitochondrial ATP production in an abundance of complex I & II substrates. *=p<0.05, ***=p<0.001 and ****=p<0.0001. **B** Direct assessment of complex I activity. *=p<0.05. **C** Direct assessment of complex II activity. *=p<0.05. **D** Direct assessment of complex IV activity **E** Astrocyte mitochondrial reactive oxygen species production assessment. *=p<0.05 **=p<0.01 In all experiments AD astrocytes are compared to relative controls using t-tests. Each experiment included 7 sAD controls, 5 sAD lines, 3 fAD control lines and 3 fAD lines. Data was analysed after at least 3 technical repeats and after 3 biological repeats were performed in each experiment.

In both sporadic and familial astrocytes a reduction in the OCR/ECAR ratio was seen (sporadic reduction 40% SD16.8%, p=0.0368, familial 27.5% reduction SD9.3%, p= 0.041). This may reflect the fact that both types of AD astrocytes appear to have a deficit in OxPHOS, but could also suggest a glycolysis deficit. (Figure 4I).

### Sporadic and Familial AD astrocyte mitochondria have a complex array of dysfunctions including increased mitochondrial ROS production, reduced complex I linked ATP production and increased complex I and II activity

*A*s both sporadic and familial astrocytes have a deficit in respiration suggesting deficiencies with OxPHOS, we measured the ATP produced linked to specific ETC complexes. Both sporadic and familial astrocytes showed a significant reduction in mitochondrial ATP production when supplied with complex I substrates (sAD: 34% reduction, p=0.046, SD 15.3%, fAD: 56% SD20.1% reduction, p=0.0485) (Figure 4A). No differences in mitochondrial ATP production were seen when complex II substrates were supplied to the permeabilised cells, suggesting no deficit in complex II activity or subsequent complexes in the respiratory chain (Figure 4A). Interestingly when specific complex I activity was measured it was shown to be significantly higher in sporadic AD astrocytes (43.4% increase, SD13.1% p=0.0078, Figure 4B), with no difference in seen in familial AD complex I activity when compared to controls (Figure 4B). When complex II activity was measured directly, sporadic AD astrocytes had significantly increased complex II activity (increase 75%, SD 21%, p=0.0061) with no difference seen in familial AD complex II activity (Figure 4C). No change in complex IV activity was seen in either the sAD or fAD astrocytes (Figure 4D). Finally, to assess the efficiency of the mitochondrial ETC we measured mitochondrial reactive oxygen species (ROS) levels in all astrocyte lines. A significant increase in ROS production was seen in both sporadic (71% increase, SD 38%, p=0.001) and familial (45% increase, SD 45%, P=0.035) AD astrocytes when compared to controls (Figure 4E). Taken together these data suggest both complex I and II have the capacity to be more active with increased Vmax measurements, however they are unable to maintain OXPHOS function and ATP levels when taken as a whole respiratory chain, possibly via substrate generation and supply; this results in higher ROS production which points towards an overall less efficient system.

### Astrocyte glycolytic function is impaired in AD astrocytes

As reported earlier in the results section, glycolysis is the main metabolic pathway used by astrocytes to maintain total cellular ATP levels. Therefore, glucose metabolism was assessed in the AD astrocytes. Figures 5A&B show the glycolysis stress test trace performed in sporadic and familial AD astrocyte lines, and how they compare to controls (Figure 5A&B). A significant reduction in glycolysis rate were seen in fAD astrocytes (78.6% reduction, p= 0.0022), with a trend towards a reduction seen in sAD astrocytes (28.2% reduction, p= 0.1124) when compared to controls (Figure 5C). A significant reduction in glycolytic capacity was seen in the sAD (40.9% reduction, p=0.0466) and fAD astrocytes (70.6% reduction, p=0.0263) (Figure 5D). Glycolytic reserve was significantly reduced in both sporadic (43% decrease, p=0.043) and familial (68% decrease, p=0.044) AD astrocytes when compared to controls (Figure 5E). No significant difference in non-glycolytic acidosis was seen in both familial and sporadic AD astrocytes when compared to controls (Figure 5.5F). Together this data suggests that both sporadic and familial AD astrocytes have impairments in their ability to metabolise glucose when compared to controls.

**Figure 5.**
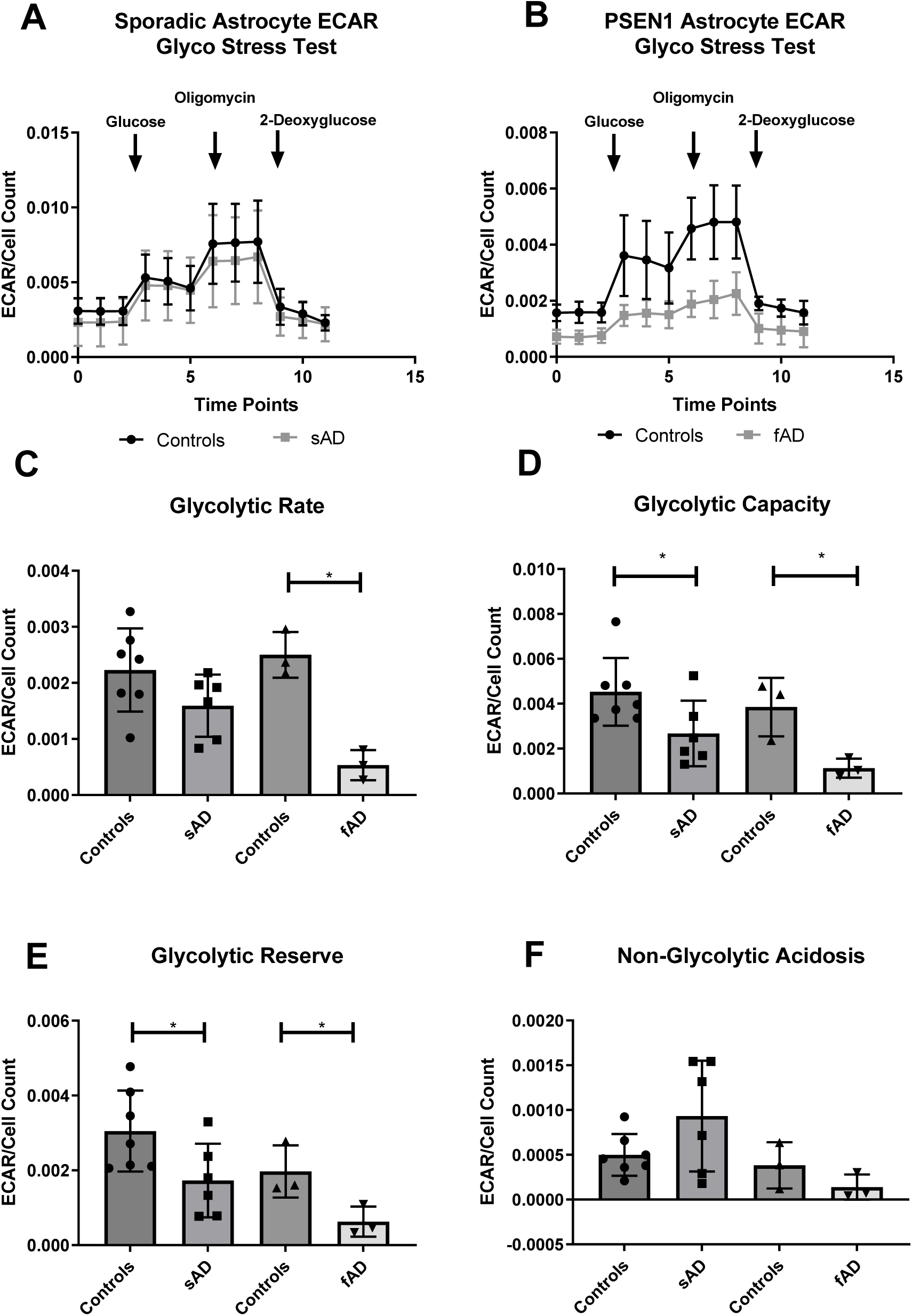
Astrocyte glycolysis assessment. **A** ECAR trace Sporadic astrocytes **B** ECAR Trace Familial Astrocytes. **C** *Glycolysis rate* *=p< 0.05. **D** Glycolytic Capacity. *=p< 0.05. **E** Glycolytic Reserve. *=p< 0.05. **F** Non-glycolytic acidosis in all experiments AD astrocytes are compared to relative controls using t-tests. Each experiment included 7 sAD controls, 5 sAD lines, 3 fAD control lines and 3 fAD lines. Data was analysed after at least 3 technical repeats and after 3 biological repeats were performed in each experiment.

### Sporadic and Familial AD astrocytes have impaired glycolytic function at multiple stages in the glycolytic pathway

The striking impairment of glycolysis in the AD astrocytes led us to investigate upstream of glycolysis, glucose uptake rates. This was measured using 2-deoxyglucose and measuring the amount of astrocyte glucose transporters. A trend to a reduction in 2-deoxyglucose uptake was seen in both sporadic (55% reduction, p=0.197) and familial (50% reduction, p=0.1422) AD astrocytes, but this was not significant (Figure 6A). Glucose transporter protein expression (GLUT1,2,&4) where all assessed via western blot. Astrocytes are known to express the GLUT1 receptor, but pervious work has also suggested that they express the GLUT 2&4 isoforms[85–87]. GLUT1 receptor protein expression was shown to be reduced in both sporadic (45% reduction, SD12%, p=0.004) and familial (39% reduction, SD 10%, p=0.023) AD astrocytes when compared to controls (Figure 6B see Figure 6C for representative blot). We could not identify expression of GLUT2 or 4 receptors in any astrocyte type using western blot analysis. To further investigate the glycolytic dysfunction, we sought to assess both the levels and activity of hexokinase which is the first key regulatory enzyme in the glycolytic pathway and importantly links glycolysis and OXPHOS. Hexokinase activity was significantly reduced in both sporadic (26% reduction SD 5.8%, p=0.001) and familial (53% reduction, SD 17%, p=0.040) astrocytes when compared to controls (Figure 6D). Hexokinase protein expression identified using western blot analysis revealed reduced expression of the protein in sporadic (33% reduction SD10.7%, P=0.011) and familial (21.5% reduction, SD 7%, P=0.044) AD astrocytes (Figure 6E, see Figure 6F for representative blot). Hexokinase mRNA expression was also assessed using qPCR which again showed a reduction in Hexokinase mRNA expression in both sporadic (40% reduction, SD 15%, p=0.022) and familial (54% reduction, SD 8.5%, p=0.0032) AD astrocytes. These results suggest a reduction in the expression and activity of the hexokinase enzyme which is central to glycolytic function and therefore could explain the glycolytic deficits seen in these cells.

**Figure 6.**
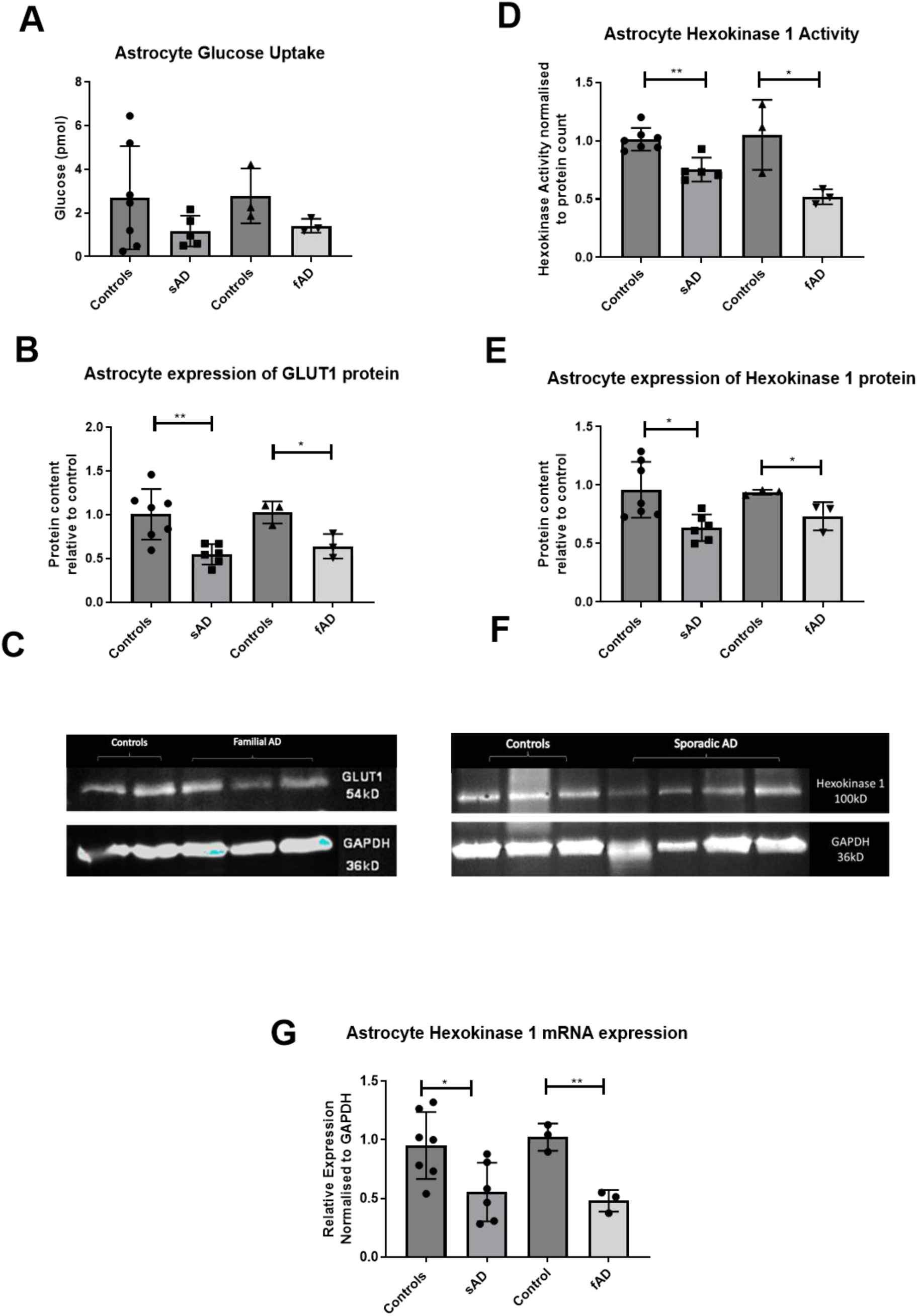
Astrocyte Glucose uptake, Hexokinase expression and activity, **A** Figure 6A Astrocyte 2-deoxyglucose uptake. **B** GLUT1 astrocyte protein expression *=p<0.05, **=p<0.01.**C** Representative western of GLUT1 blot **D** Hexokinase 1 activity*=p<0.05, **=p<0.01. **E** Hexokinase protein expression*=p<0.05. **F** Representative blot of Hexokinase protein expression Figure 6G Hexokinase mRNA expression. In all experiments AD astrocytes are compared to relative controls using t-tests. Each experiment included 7 sAD controls, 5 sAD lines, 3 fAD control lines and 3 fAD lines. Data was analysed after at least 3 technical repeats and after 3 biological repeats were performed in each experiment.

### Astrocyte hexokinase activity correlates with markers of astrocyte metabolic reserve and metabolic output

To assess the significance of hexokinase deficits in astrocytes we next correlated several markers of astrocyte metabolic reserve, metabolic output and mitochondrial function highlighted as deficient earlier on in the results section with hexokinase enzyme activity. Hexokinase activity correlated positively with both MSRC (R=0.338, p=0.047, Figure 7A) and glycolytic reserve (R=0.490, p=0.011, Figure 7B). With regards to astrocyte metabolic outputs, astrocyte hexokinase activity correlated positively with extracellular lactate (R=0.434, p=0.012, Figure 7C) and total cellular ATP (R=0.500, p=0.01, Figure 7D). Mitochondria ROS level correlated negatively with hexokinase activity (R=0.451, p=0.016, Figure 7E) and there was no significant correlation between hexokinase activity and MMP (R=0.229, p=0.139, Figure 7F). These results suggest that several of the deficits seen in astrocyte metabolic function may be interdependent on the activity of astrocyte hexokinase 1.

**Figure 7.**
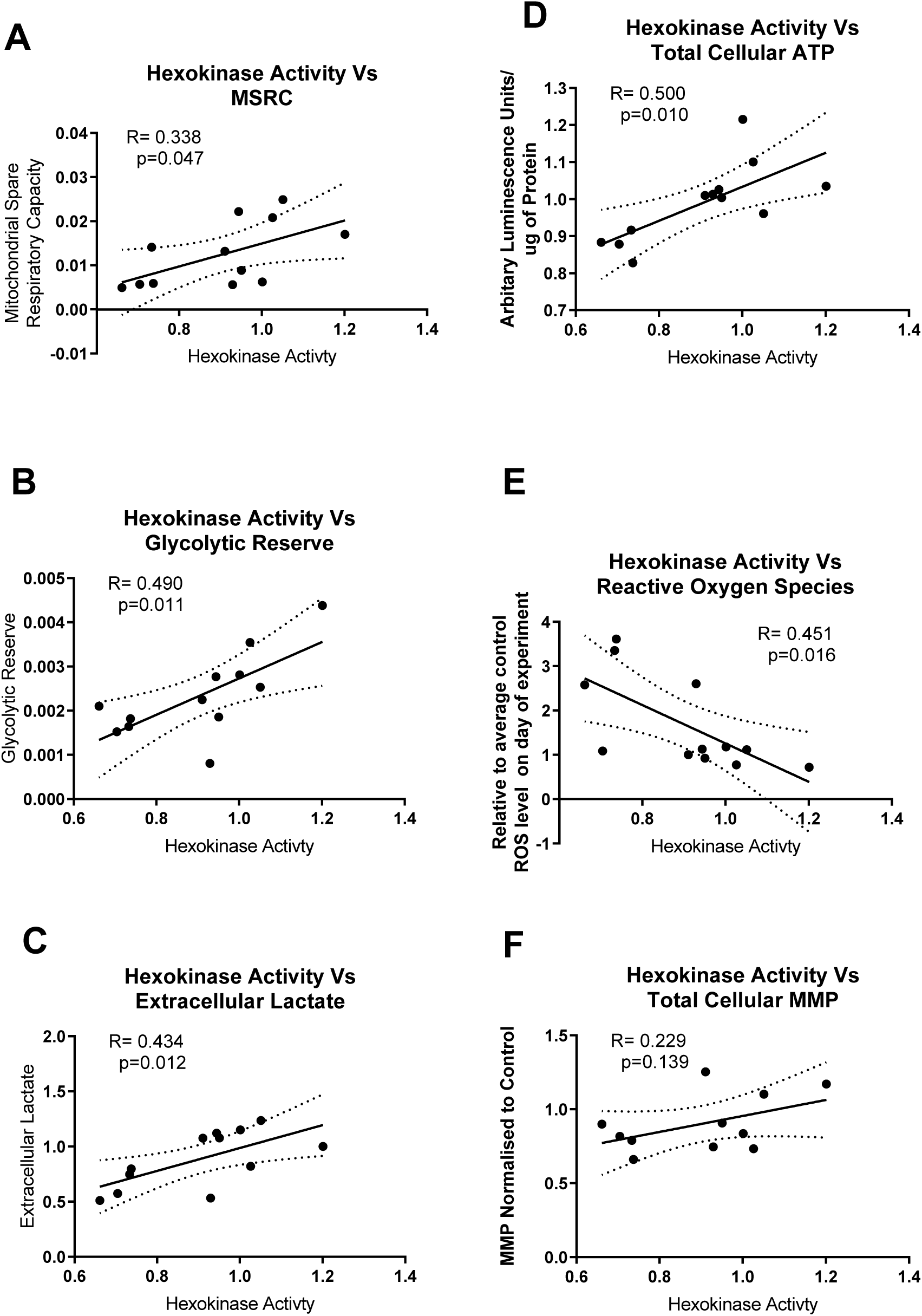
Astrocyte Hexokinase activity correlates with metabolic deficits seen in glycolysis and mitochondrial function. **A** Hexokinase vs MSRC correlation **B** Hexokinase vs Glycolytic reserve **C** Hexokinase vs Extracellular lactate **D** Hexokinase vs Total Cellular ATP **E** Hexokinase vs ROS **F** Hexokinase vs MMP. In each correlation 7 controls and 5 sAD cell lines are plotted. The correlations are performed using a 2-tailed p-value and a Pearson correlation coefficient.

### Restoring hexokinase expression improves some markers of astrocyte metabolic output, reduces mitochondria ROS, but does not improve mitochondrial morphological changes in sporadic AD Astrocytes

Next, we assessed if correcting the deficit in astrocyte hexokinase expression would improve astrocyte metabolic output. We picked the 3 sAD astrocyte lines with the largest hexokinase deficit and the 3 fAD lines to transduce with an AVV containing the hexokinase 1 gene. Figure 8A shows the increased expression of the hexokinase protein in the astrocyte cells after transduction with the hexokinase 1 containing AVV. Expression of the hexokinase protein increased by 329% in sporadic controls, p<0.0001, 216% in sAD astrocytes, p=0.05, 184% in familial control astrocytes p=0.05, and 210% in fAD astrocyte lines, p=0.035. Figures 8B-C show example westerns for both sporadic and familial AD hexokinase overexpression. In sAD astrocyte lines transduction with the hexokinase 1 containing AVV led to a significant increase in total cellular ATP (21.6% increase, SD 7.3%, P=0.041, Figure 8D), a significant increase in extracellular lactate (41.1% increase, SD 11.4%, p=0.027, Figure 8E) and a significant reduction in mitochondrial ROS (49.5% decrease, SD 14.3%, P=0.026, Figure 8F) when compared to controls. Interestingly, in fAD astrocytes, transduction with the hexokinase 1 containing AAV did not increase total cellular ATP (Figure 8G), nor increased extracellular lactate (Figure 8H), or lead to a significant reduction in mitochondrial ROS (Figure 8I). Abnormal mitochondrial morphological parameters, abnormal in sAD and fAD astrocytes were not improved with AVV transduction (Supplementary figure 1). This data suggests that in sporadic AD astrocytes the deficits in glycolysis and mitochondrial ROS can be corrected by increasing hexokinase 1 expression but has no effect on the mitochondrial morphological abnormalities seen. Interestingly, this striking restoration does not occur in the fAD astrocytes, suggesting the mechanism by which those deficits occur are different between sAD and fAD astrocytes.

**Figure 8.**
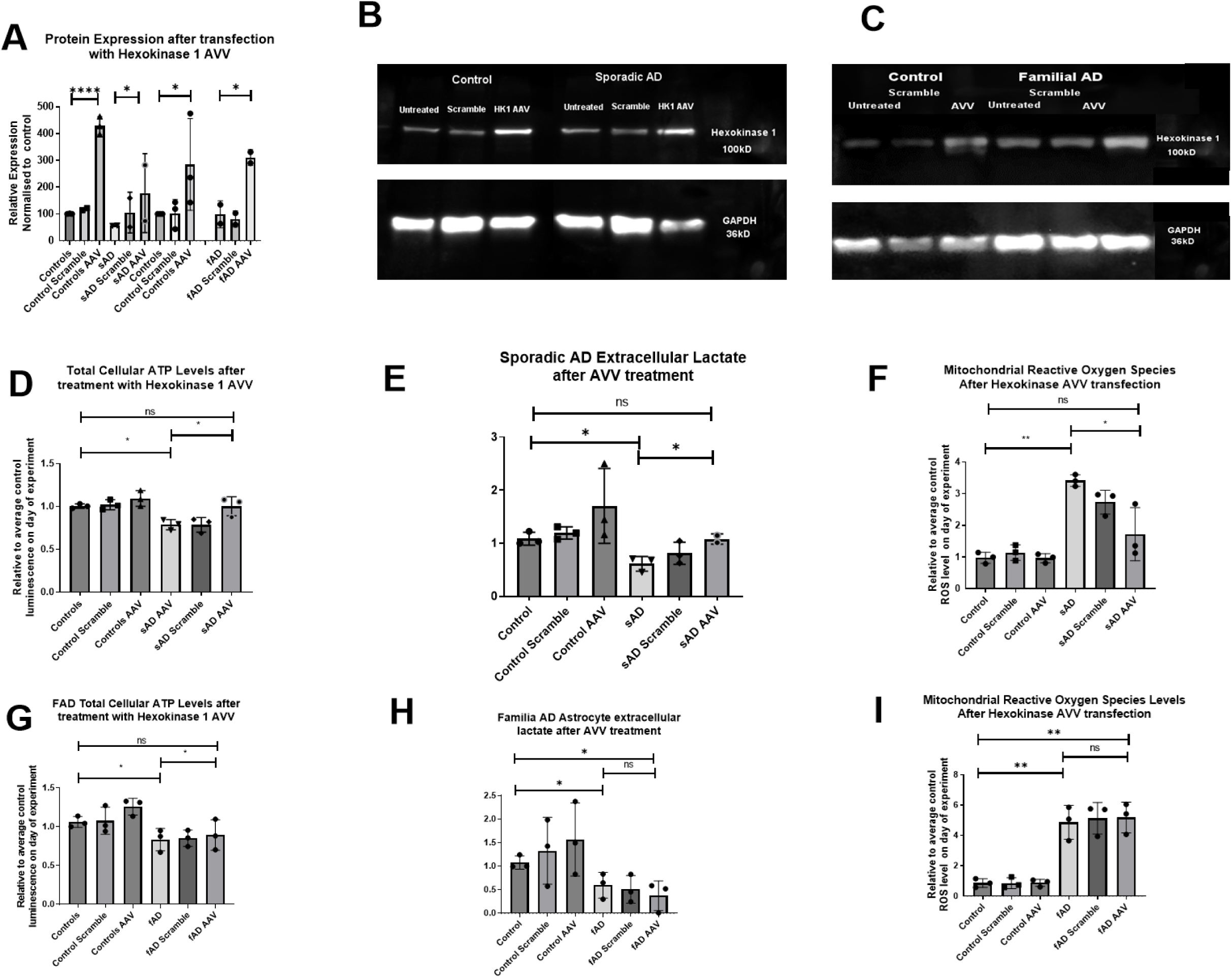
Transfection with an hexokinase containing AAV vector restores metabolic outputs in sAD astrocytes but not fAD. **A** Hexokinase protein expression after transfection with the hexokinase containing AAV. *=p<0.05, ****= p<0.0001. **B** Representative western for hexokinase expression in sAD astrocytes and controls. **C** Representative western for hexokinase expression in fAD astrocytes and controls. **D** Total cellular ATP levels after AAV transfection in Sad. *=p<0.05 **E** Extracellular lactate levels after AAV transfection in sAD lines. *=p<0.05 **F** Astrocyte mitochondrial ROS production after AAV transfection sAD lines. *=p<0.05, and **=p<0.01. **G** Total cellular ATP levels after AAV transfection in fAD. *=p<0.05 **H** Extracellular lactate levels after AAV transfection in sAD lines. *=p<0.05 **I** Astrocyte mitochondrial ROS production after AAV transfection fAD lines. **=p<0.01. In all experiments AD astrocytes are compared to relative controls using t-tests. Each experiment included 3 sAD controls, 3 sAD lines, 3 fAD control lines and 3 fAD lines. Data was analysed after at least 3 technical repeats and after 3 biological repeats were performed in each experiment.

### Astrocyte metabolic output and capacity markers correlate with neuropsychological markers in sporadic AD astrocytes but not hexokinase 1 activity

In our previous paper we investigated if MSRC and MMP measured in sAD fibroblasts correlated with neuropsychological markers of AD [88]. As we have now identified a distinct metabolic phenotype in sAD astrocytes to that identified in the fibroblasts from which the astrocytes are derived, we decided to investigate if the same correlations were present. Although similarities are seen in the mitochondrial and glycolytic deficits in both sAD fibroblasts and astrocytes, sAD fibroblasts have more pronounced structural change to their mitochondrial network when compared to sAD astrocytes. This includes an increase in perinuclear mitochondria, and a reduced mitochondrial number that is not seen in astrocytes. Astrocytes have a more profound metabolic deficit signified by reductions in total cellular ATP and glycolytic rate, which is not seen in sAD fibroblasts. Astrocytic glycolytic function is more significantly impaired than in fibroblasts for sAD patient derived cells [88], therefore we also investigated if glycolytic reserve and extracellular lactate correlated with neuropsychological markers of AD. Neuropsychological data was only available for sAD astrocytes and their control lines, therefore fAD astrocytes could not be assessed. Semantic fluency, immediate and delayed recall were assessed as these were the neuropsychological markers previously investigated, and are known to be affected in the early stages of AD. A significant positive correlation was seen with immediate recall and MSRC (R=0.669, p=0.005), (Supplementary Figure 2B). A positive correlation, which did not reach significance was seen between MSRC and semantic fluency (Supplementary Figure 2A) and delayed episodic recall (Supplementary Figure 2C). Positive correlations that were not significant were seen between MMP and semantic fluency, (Supplementary 2D), immediate recall, (Supplementary Figure 2E) and delayed episodic recall (Supplementary Figure 2F). Glycolytic reserve showed a trend towards a positive correlation with semantic memory (R=0.524, p=0.054), (Supplementary Figure 3A). A significant positive correlation was seen between both glycolytic reserve and immediate recall (R=0.570, p=0.033) (Supplementary Figure 3B), and glycolytic reserve and delayed recall (R=0.541, p=0.045) (Supplementary Figure 3C). Extracellular lactate showed significant positive correlations with all three neuropsychological tests, semantic fluency (R=0.634, p=0.014, Supplementary Figure 3D), Immediate recall (R=0.617, p=0.018, Supplementary Figure 3E), delayed recall (R= 0.578, p=0.038, Supplementary Figure 3F). After identifying the above correlations, as we had done previously in our paper using fibroblasts from the same participants we assessed the correlations after controlling for brain reserve, length of education and participant age [88]. Only correlations that were statistically significant were assessed. After controlling for the 3 factors, positive correlations remained between immediate recall and glycolytic reserve (r=0.824, p=0.003), delayed recall and glycolytic reserve (r=0.658, p=0.039) and semantic fluency and extracellular lactate levels (r=0.865, p=0.003). A trend towards significance was seen with the correlation between immediate recall and MSRC (r=0.586, p=0.075), but the correlations between extracellular lactate and immediate recall (r=0.552, p=0.123) and delayed recall were no longer significant (r=0.368, p=0.330). Supplementary Table 3 displays these correlations. Finally, as hexokinase 1 activity correlates with significantly with the above-mentioned metabolic deficits seen in AD astrocytes, we investigated in hexokinase 1 activity correlated with the aforementioned neuropsychological markers. A positive correlation was seen between immediate recall (r= 0.269, p=0.0836, Supplementary figure 4A,) and Delayed recall (r=0.318, p=0.055, Supplementary figure 4B) but was not significant. Semantic fluency did show a significant positive correlation with hexokinase 1 activity (r=0.354, p=0.043, supplementary figure 4C). In conclusion this data highlights that both mitochondrial and glycolytic astrocytic parameters shown to be abnormal in AD correlate with neuropsychological tests known to be affected early in the course of the condition.

## Discussion

In this study we have shown that astrocytes derived from patients with both sporadic and familial AD have deficits in key metabolic pathways, including several mitochondrial abnormalities and glycolytic abnormalities. These abnormalities are strikingly similar between sAD and fAD astrocytes. We show that some of these abnormalities correlate with neuropsychological abnormalities seen early AD. Hexokinase 1 is a key enzyme in the glycolytic pathway which links the major metabolism pathways of glycolysis and OXPHOS. We find that overexpression of hexokinase 1 remarkably restores most metabolic abnormalities in the sAD astrocytes, suggesting that in sAD hexokinase 1 activity is a therapeutic target to restore astrocyte glucose metabolism, which may have an effect on cognitive performance.

### The AD astrocyte metabolic phenotype

Our work builds upon limited studies available to date investigating metabolism in astrocytes derived from AD patients. One of the few published studies utilizing astrocytes derived from AD patients, found a decreased rate of glycolysis, decreased glycolytic reserve and lower extracellular lactate levels in astrocytes derived from 3 PSEN1 mutation carriers when compared to controls [89]. In a further study using iPSC derived astrocytes from participants with sporadic forms of AD decreased expression of hexokinase, and reduced glucose uptake was also seen [90]. In this study we see similar changes in glycolytic pathways in astrocytes derived via an alternative reprogramming method in both a sporadic and PSEN1 familial AD cohort; furthermore, we add to the literature with the finding that correcting the hexokinase expression deficit in sporadic AD can correct several of the glycolytic deficits seen in these astrocytes, highlighting a potential therapeutic target.

Other studies that have investigated glycolysis in AD astrocytes have used cells from either post-mortem (PM) brain tissue, or animal models of disease, although specific data about astrocyte glycolysis is limited. The activity of Phosphofructokinase (PFK), the enzyme that converts fructose-6-phosphate into fructose-1,6-bisphosphate is thought to have increased activity in the astrocytes of people with AD [91]. There is also evidence that several other enzymes of the glycolytic pathway are upregulated in the PM AD brain including lactate dehydrogenase and pyruvate kinase [92, 93], although astrocytes are not specifically targeted in these studies. In our study, which focuses on people at an early stage of AD, we have shown parameters of the overall rate of glycolysis are lower in AD astrocytes and that hexokinase 1 activity and expression is reduced in both sporadic and familial AD astrocytes.

The results from our study and the evidence of increased enzyme activity could be explained by a reduced glucose uptake by the astrocyte. There is evidence that the glucose transporters GLUT1&3 have reduced expression as AD progresses [90, 94], and we have found a reduction in the expression of the GLUT1 transporter in both sAD and fAD. This could lead to a situation in which astrocytes have lower overall rates of glycolysis but increase enzyme activity to compensate. If hexokinase 1 activity is reduced this may also contribute to an increase in the activities of the other glycolytic enzymes. Alternatively, a reduction in astrocyte glycolysis may lead to increased neuronal glycolysis, which may explain why increased enzyme content is seen in AD PM specimens. We did not measure the other glycolytic enzymes as part of this project, but this would be important work for future projects to focus on. Human imaging studies also suggest that the brain has a reduced glucose uptake in both ageing and AD [95] this could be explained by a reduced number of glucose receptors in the brain, or a reduction in astrocyte hexokinase as seen in our study.

Studies of enzyme activity in fibroblasts from patients with sAD have shown a reduction in the activity of hexokinase, the first enzyme in the glycolytic pathway, [96]. It has also been shown that fibroblasts from sAD patients rely more on glycolysis than controls but have a limited ability to increase glucose uptake [97]. We and others have identified similar deficits in the astrocytes derived from patient fibroblasts[90]. It could be argued that the changes in glycolytic function seen in this study merely reflect changes in fibroblast glycolysis and not primary astrocytic glycolytic defects. The metabolic profile of the astrocytes produced in this project is different to that of the fibroblasts that they are reprogrammed from. The parent fibroblasts do not have deficits in glycolytic capacity or glycolysis rate and on a group level total cellular ATP is affected much less [24, 25]. Furthermore, data highlighting that the brain becomes less sensitive to glucose uptake through the course of AD, and the fact that the majority of glucose metabolism within the brain is thought to be linked to astrocytic maintenance of glutamatergic synapses [98], this points towards a deficit in astrocyte glucose metabolism in human studies. Separate cohorts of astrocytes reprogrammed using different technology and from different patient groups have shown abnormalities astrocyte glycolysis [89] [90] corroborating the changes to glycolysis in primary astrocytes seen in this study not being tessellations of fibroblast glycolytic function.

Astrocytes can shuttle lactate to neurons as a fuel source, with neurons having a preference for lactate over glucose in times of increased energy expenditure [99–101]. In this study we have shown that astrocytes from both sporadic and familial AD patients have reduced extracellular lactate levels, which has potential consequences for the neuron astrocyte relationship. Neurons rely mainly on OxPHOS to meet their energy requirements [49], as a result of this they are at high risk of oxidative damage from free radicals produced via the electron transport chain. It has been shown that lactate preference, as an energy source, in neurons allows for the utilization of neuronal glucose to produce antioxidant molecules such as glutathione via the pentose phosphate pathway (PPP) [101]. If the supply of astrocyte lactate to neurons is impaired, this could lead to an increase in oxidative damage in AD neurons due to glucose being diverted away from the PPP. Oxidative damage to neurons has been reported when AD astrocytes are co-cultured with non-AD neurons [89], and in multiple PM studies of the AD brain [102–104]. We have also shown in this study that sporadic and familial AD astrocytes have higher levels of mitochondrial ROS when compared to controls. ROS are used as a signalling molecule and can be released by astrocytes. The higher levels of ROS seen in AD astrocyte will contribute to dysfunction of the astrocyte itself but may also contribute to oxidative damage of neurons seen in PM brains and co-culture studies. This suggests that decreased release of lactate or the increased release of ROS from astrocytes, may have a role in the pathogenesis of AD, and therefore would make a suitable future therapeutic target.

The reduction in astrocyte total cellular ATP reported in this study is likely to also affect the astrocyte-neuron relationship. ATP is used as a signalling molecule within the brain, and astrocyte released ATP is known to modulate synaptic activity [105–107]. It is thought that astrocyte ATP in part can alter the survivability of a forming synapse via its actions on the P2Y receptor[108]. As synaptic loss is an early sign in AD, support of astrocyte metabolism to increase total cellular ATP may help to prevent this.

We also report changes in mitochondrial function and morphology. Previous work on human derived fAD astrocytes has suggested that the oxygen consumption rate (OCR) is higher than that of matched controls [89]. Trends towards deficits in MSRC have also been identified but have not been shown to be significant [109]. Both these studies focus on the same 3 PSEN1 astrocyte lines. The changes in MSRC are consistent with what we found in this study although, we have shown the MSRC deficit to be significant. The basal OCR in our study was significantly lower in AD astrocytes which is the opposite finding seen by Oksanen and colleagues [89]. The difference seen here may be explained by the fact that we have studied point mutations in the PSEN1 gene that cause AD, whereas the Oksanen group have studied astrocytes created from an exon 9 deletion model of AD. In this study we have shown that astrocytes are more dependent on glycolysis than OxPHOS for ATP production. Potentially the exon 9 deletion causes a more severe glycolysis deficit than the point mutations that we have investigated in this study, This would lead to a greater reliance on OxPHOS, and my lead to higher OCR rates.

The morphology of the mitochondrial network is altered in both sporadic and familial AD astrocytes showing longer mitochondria that are more interconnected. Potentially these changes could suggest that the astrocyte is under metabolic stress, and the trend towards an increase in the percentage of perinuclear mitochondria would suggest a collapsing mitochondrial network. The identified increased mitochondrial ROS levels in AD astrocytes may also point towards mitochondrial stress and inefficiency. As the AD astrocytes have a deficit in glycolysis as well as OxPHOS, it is difficult to know if the mitochondrial changes seen are a consequence of glucose metabolism failure, leading to an increased need for OxPHOS, or if the astrocytes have inherent deficits in mitochondrial function. The fact that MSRC and glycolytic reserve were significantly reduced in both sporadic and familial AD astrocytes, suggests primary deficits in both metabolic pathways, as opposed to the glycolysis defect causing a secondary mitochondrial functional change. The reduction in mitochondrial cellular ATP production when complex I substrates are supplied to astrocytes, suggests a primary deficit in OxPHOS, but when we investigated the function of complex I, II, and IV directly no deficit in activity was identified. Interestingly, in sporadic AD astrocytes the activity of both complexes I and II was significantly higher than comparative controls, which may in part help to explain the increased ROS level seen in AD astrocytes. It has been previously shown that increase mitochondrial ROS levels can stimulate both glucose uptake and GLUT1 expression in myoblasts [110]. The increased ROS levels we see in this study may be a compensatory mechanism by the astrocyte ETC to try and increase glucose uptake. The deficits in the activity and expression of hexokinase could explain both the deficits in glycolysis and OxPHOS. Reduced hexokinase activity would lead to a reduction in overall glycolysis, this would reduce substrates available for ETC and hence effect the maximum mitochondrial oxygen consumptions and therefore MSRC.

### Correction of hexokinase deficit

As with previous studies that have investigated glycolytic and mitochondrial changes in AD patient derived astrocytes, we have identified several deficits in both glycolysis and mitochondria structure and function. We investigated what might be the underlying cause of these changes in glycolytic function and mitochondrial structure and function in AD astrocytes by correcting the deficit in hexokinase activity and expression using an AVV. We chose this enzyme as it is the first enzymatic step in the glycolysis pathway, but also is positioned within the mitochondrial outer membrane. Its function has been shown to affect mitochondrial ETC activity, and hence deficits in the enzyme expression and function may explain changes to both glycolysis and mitochondrial structure and function. By increasing the expression of hexokinase 1 we were able to ameliorate the deficits in total cellular ATP and extracellular lactate levels in sporadic AD astrocytes. Transduction with the hexokinase 1 AVV also reduced the mitochondrial ROS levels. This suggests that increasing hexokinase 1 expression recovers the function of the glycolytic pathway allowing restoration of total cellular ATP and extracellular lactate levels. As discussed above, correcting these abnormalities is likely to have significant benefits to the neuron-astrocyte relationship and also synaptic maintenance. The reduction in mitochondrial ROS seen also suggests that an element of the astrocyte mitochondrial dysfunction is secondary to, or dependent on, AD glycolysis not proving enough substrates for the ETC.

In sporadic AD astrocytes increasing hexokinase 1 expression did not correct any of the mitochondrial structural abnormalities seen. It could be that longer exposure to increased levels of hexokinase 1 is needed before the astrocyte mitochondria remodel to a more normal morphology. Potentially the mitochondrial structural changes seen in AD astrocytes are driven by another pathology. We have shown in sporadic and familial AD fibroblasts that Drp1, a mitochondrial fission protein, has reduced expression [24]. The same abnormality could be present in AD astrocytes and maybe the driver of mitochondrial structural change.

Increasing the expression of hexokinase 1 in fAD astrocytes did not improve any of the markers of glycolytic function or mitochondrial structure and function. The fAD astrocytes had a greater deficit than sAD astrocytes in most mitochondrial and glycolytic measurements. Potentially the increased expression of hexokinase in fAD was not significant enough to overcome the deficits seen in glycolysis. It could also be suggested that deficits in fAD astrocyte glycolysis and mitochondrial function are driven by a separate mechanism, possibly the metabolic phenotype is driven via a mitochondrial mechanism. In this study we were not able to measure amyloid beta levels, but it is likely the fAD astrocyte expresses higher levels of this protein which is known to impede both glycolysis and mitochondrial function.

### Astrocyte metabolism correlations with neuropsychological changes

In our previous paper we have shown that deficits in MSRC and MMP correlate with neuropsychological changes seen early in AD. We repeated this analysis with the patient derived sporadic astrocytes and saw very similar correlations. As well as looking at mitochondrial functional parameters we also investigated if glycolytic reserve, and extracellular lactate correlated with neuropsychological changes. We found in both cases that immediate and delayed recall correlated significantly with extracellular lactate levels and glycolytic reserve. Extracellular lactate also correlated with semantic fluency significantly. When studying fibroblasts, significant correlations were not seen with glycolysis parameters. This could reflect the fact that the astrocyte depends on glycolysis to a higher degree than the fibroblast. It is also interesting that extracellular lactate correlates with neuropsychological measures considering the importance lactate has in neuronal metabolism. The correlations between extracellular lactate and immediate and delayed recall did not survive controlling for factors that can affect scoring on neuropsychological tests such as age, brain reserve and years of education. This may be explained by the small sample size of study. The fact that both MSRC and glycolytic reserve correlate with neuropsychological measures is interesting, as both these tests assess capacity in metabolic systems. We and many other groups have shown that metabolic capacity is affected in AD [23-25, 29, 111, 112]. We also investigated if astrocyte hexokinase activity correlated with the same neuropsychological scores. Positive correlations were seen in each comparison with hexokinase activity, but only delayed recall showed significance. The stronger correlations between extracellular lactate and glycolytic reserve may suggest that the factors that affect the metabolic relationship between the neuron and astrocyte are more important for cognition then the cause of the metabolic deficit.

This is the first study to show that astrocytic glycolytic reserve, extracellular lactate and MSRC correlates with scoring on neuropsychological tests shown to be affected early in AD. This is an important finding, as there is potential to develop a metabolic biomarker from these functional deficits. The development of clinical studies investigating astrocyte metabolism enhancers in AD could also use these correlations to track compound effect. Caution in interpreting this correlation is needed as this is a small cohort of sporadic patients. Further work on larger cohorts of sporadic AD patients would be needed before these correlations could be developed into a clinically useful tool.

This study is limited by sample size but is one of the larger studies investigating metabolic abnormalities in astrocytes derived from both sporadic and familial AD patients. Further work is needed to identify the mechanism behind metabolic dysfunction in familial AD astrocytes, and also if replacement of astrocyte hexokinase expression and activity improves the functional relationship between astrocytes and neurons in AD.

## Conclusions

In this study we show that astrocytes derived from patients with sporadic or familial AD have deficits in both mitochondrial function and glycolysis. We have shown that overexpressing hexokinase 1 can correct several of the glycolytic deficits in sporadic AD astrocytes but not familial. These deficits correlate with neuropsychological tests which show early change in AD. The metabolic deficits could have a profound effect on the astrocytes ability to support neurons in co-culture, and are a future therapeutic target for both sporadic and familial AD.

## Abbreviations

Aβ: Amyloid beta
AD: Alzheimer’s Disease
ATP: Adenosine triphosphate
AVV: Adenovirus Viral Vector
CNS: Central nervous system
ECT: Electron transport chain
iNPC: Induced Neuronal progenitor cells
iPSC: induced Pluripotent stem cells
MMP: mitochondrial membrane potential
MOI: Multiplicity of Infection
MRSC: mitochondrial respiratory spare capacity
OxPHOS: Oxidative Phosphorylation
OCR: oxygen consumption rate
PBS: Phosphate buffer saline
PBST: Phosphate buffer saline with tween
PFK: Phosphofructokinase
PPP: Pentose Phosphate shunt Pathway
TMRM: tetramethlyrhodamine

## Declarations

All authors report no competing interests.

## Funding

This research was funded by Wellcome 4ward north (216340/Z/19/Z), ARUK Yorkshire Network Centre Small Grant Scheme (Ref: ARUK-PCRF2016A-1), Parkinson’s UK (F1301), European Union Seventh Framework Programme (FP7/2007–2013) under grant agreement no. 601055, NIHR Sheffield Biomedical Research Centre award.

## Acknowledgements

This research was supported by the Wellcome 4ward North Academy and ARUK

Yorkshire Network Centre Small Grant Scheme (to SMB), Parkinson’s UK (to HM, F1301) and the NIHR Sheffield Biomedical Research Centre. This is a summary of independent research carried out at the NIHR Sheffield Biomedical Research Centre (Translational Neuroscience). The views expressed are those of the author(s) and not necessarily those of the NHS, the NIHR or the Department of Health and Social Care (DHSC). Support of the European Union Seventh Framework Programme (FP7/2007–2013) under grant agreement no. 601055, VPH-DARE@IT to AV is acknowledged. PJS is supported by an NIHR Senior Investigator award. The support of the NIHR Clinical Research Facility—Sheffield Teaching Hospital is also acknowledged.

## Notes

### Competing Interest Statement

The authors have declared no competing interest.

