## Supplemental tables and figures for "Deficits in mitochondrial function and glucose metabolism seen in sporadic and familial Alzheimer’s disease derived Astrocytes are ameliorated by increasing hexokinase 1 expression"

**Institutions**

1. **Sheffield Institute for Translational Neuroscience, University of Sheffield, 385a Glossop Rd, Broomhall, Sheffield S10 2HQ.**
2. **F11 Chemistry Building, University of Sydney, Australia**
3. **Department of Life Sciences, Brunel University London, Kingston Lane, Uxbridge, Middlesex, UB8 3PH**

### Supplementary Tables and Figures

| Antibody | Species | Dilution | Supplier |
|---|---|---|---|
| Vimentin | chicken | 1:200 | Abcam (AB5733) |
| Nestin | Mouse | 1:200 | Abcam (AB18102) |
| CD44 | Rabbit | 1:200 | Abcam (AB157107) |
| S100β | Rabbit | 1:200 | Abcam (AB868) |
| PAX6 | Rabbit | 1:1000 | Abcam (AB5790) |
| GFAP | Rabbit | 1:200 | Dako (Z0334) |
| NDGR2 | Mouse | 1:200 | Santacruz (sc-376202) |
| ALDH1L1 | Rabbit | 1:1000 | Abcam (AB87117) |
| **Secondary Antibodies** | | | |
| Alexia 488 | Rabbit | 1:1000 | Invitrogen |
| Alexia 488 | Mouse | 1:1000 | Invitrogen |
| Alexia 488 | Chicken | 1:1000 | Invitrogen |
| Alexia 568 | Rabbit | 1:1000 | Invitrogen |
| Alexia 568 | Mouse | 1:1000 | Invitrogen |
| **Western Primary Antibodies** | | | |
| GLUT1 | Rabbit | 1:2000 | Proteintech (66290-1-Ig) |
| GLUT2 | Rabbit | 1:250,500,1000 | Abcam (ab54460) |
| GLUT4 | Rabbit | 1:250,500,1000 | Abcam (ab33780) |
| Hexokinase 1 | Rabbit | 1:2000 | Abcam (ab648) |
| GAPDH | Mouse | 1:2000 | Proteintech (60004-1-0g) |
| **Secondary Western Antibodies** | | | |
| | Rabbit | 1:5000 | |
| | Mouse | 1:10000 | |

**Supplementary Table 1 |** ***Antibodies used for Immunochemistry and western blotting*** In this table details of antibodies used to characterise both iNPCs and astrocytes are displayed with supplier and dilution that they were applied to the cells.

| Gene | Forward | Reverse | Supplier | Cells tested |
|---|---|---|---|---|
| Hexokinase 1 (HK1) | GTCCAAGAAGTCAGAGATG CAGG | CTGCTGGTGAAAATCCGT AGTGG | Merck | Astrocytes |

**Supplementary Table 2 |** ***Hexokinase 1 gene primer sequences***

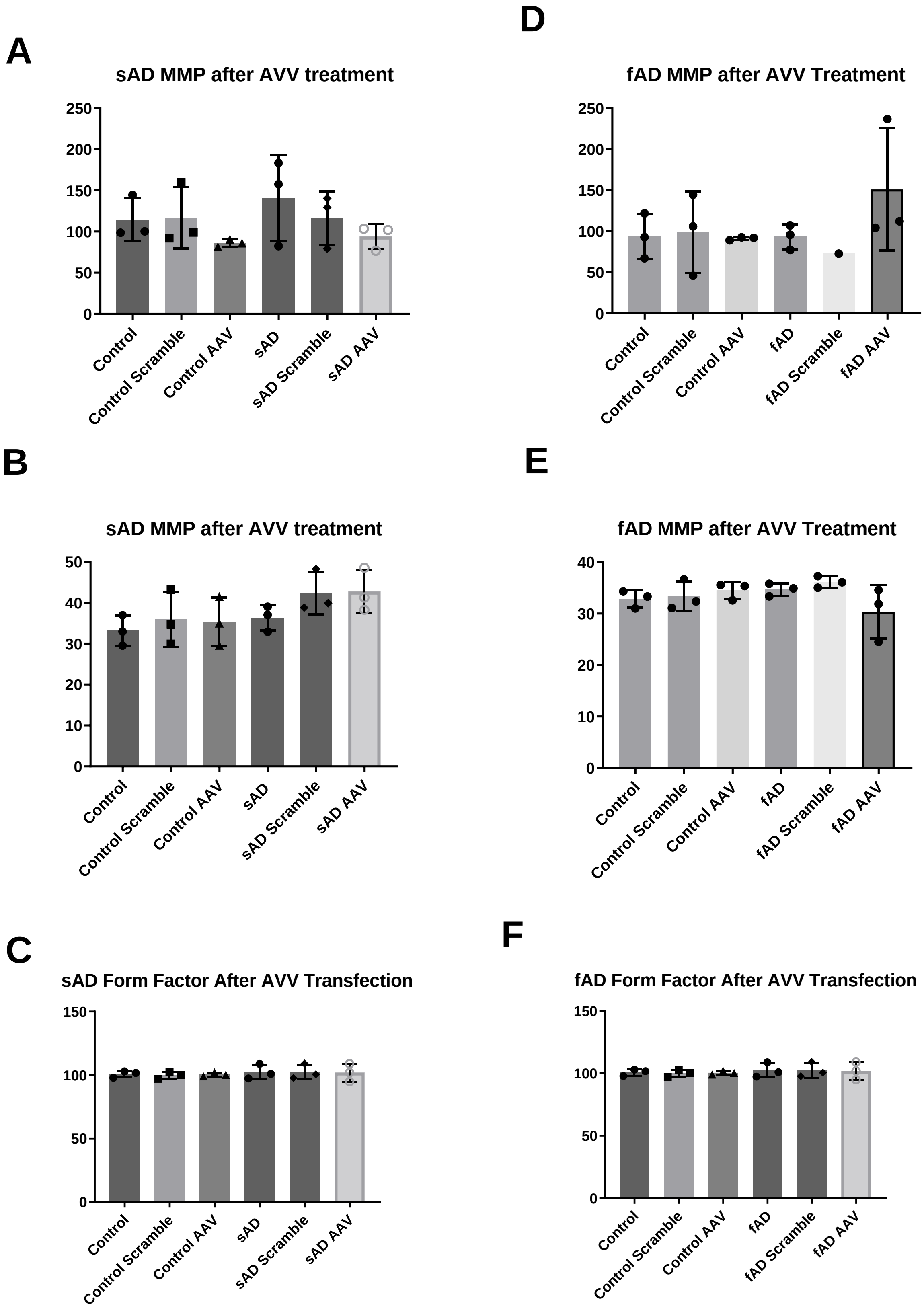
A
sAD MMP after AVV treatment
250
200
150
100
50
0
Control
Control Scramble
Control AAV
sAD
sAD Scramble
sAD AAV
D
fAD MMP after AVV Treatment
250
200
150
100
50
0
Control
Control Scramble
Control AAV
fAD
fAD Scramble
fAD AAV
B
sAD MMP after AVV treatment
50
40
30
20
10
0
Control
Control Scramble
Control AAV
sAD
sAD Scramble
sAD AAV
E
fAD MMP after AVV Treatment
40
30
20
10
0
Control
Control Scramble
Control AAV
fAD
fAD Scramble
fAD AAV
C
sAD Form Factor After AVV Transfection
150
100
50
0
Control
Control Scramble
Control AAV
sAD
sAD Scramble
sAD AAV
F
fAD Form Factor After AVV Transfection
150
100
50
0
Control
Control Scramble
Control AAV
fAD
fAD Scramble
fAD AAV

**Supplementary Figure 1 | <u>Mitochondrial morphological parameters after transfection with an AAV containing the hexokinase 1 gene.</u>** **A** sAD MMP values, **B** sAD long mitochondrial proportion. **C** sAD Form factor. **D** fAD MMP values, **E** fAD long mitochondria proportion, **F** fAD Form factor.

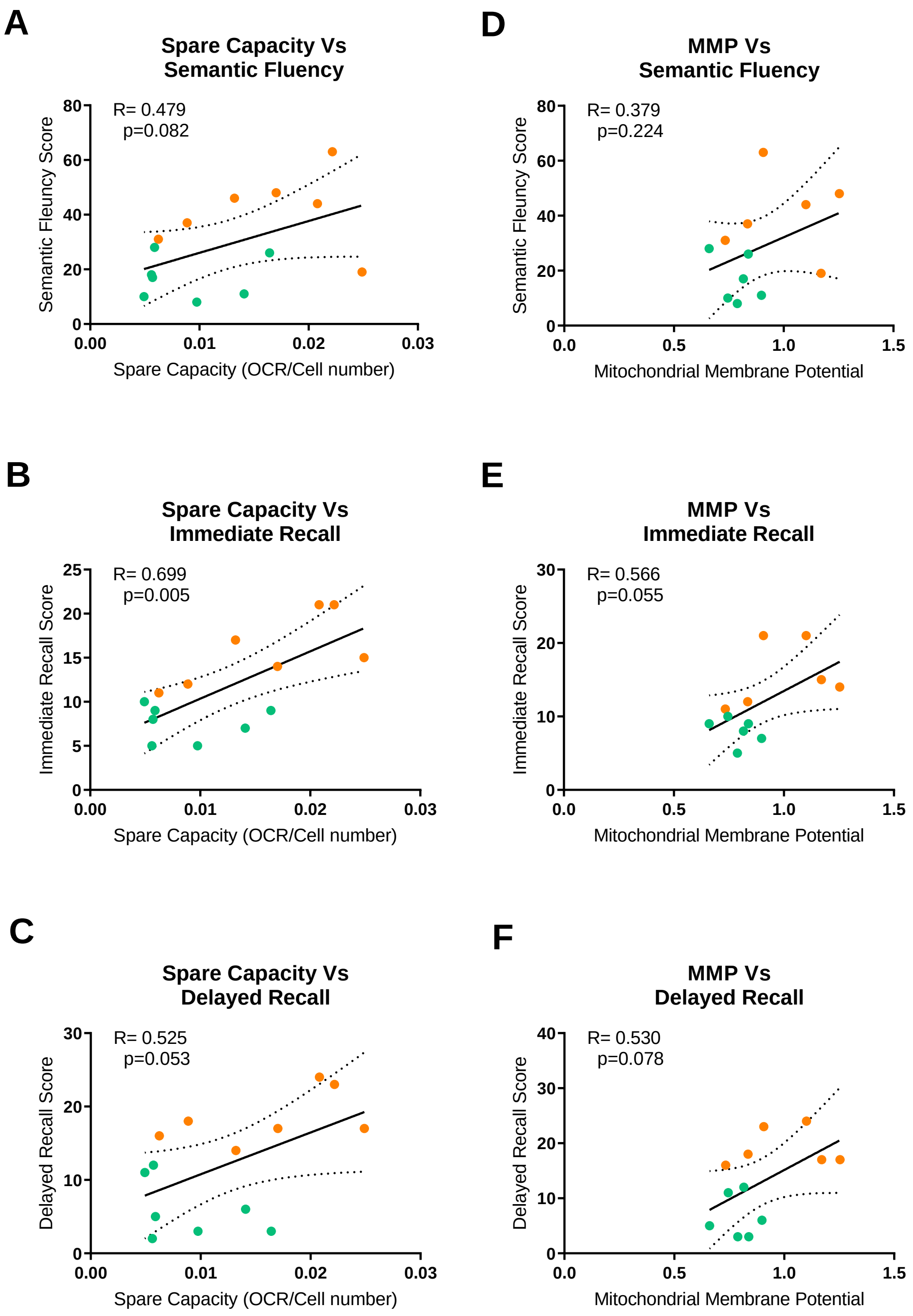


**Supplementary Figure 2 | <u>*MSRC and MMP correlations with Neuropsychological tests*</u>**

This figure highlights the mitochondrial spare respiratory capacity (MSRC) correlations with immediate recall (Figure 2A) semantic memory (Figure 2B) and delayed recall (Figure 2C). Mitochondrial membrane potential correlations with semantic memory (Figure 2D) immediate recall (Figure 2E) and delayed recall (Figure 2F) are also displayed. Green points represent sporadic AD patients and orange points represent control subjects. In each correlation 7 controls and 5 sAD cell lines are plotted. The correlations are performed using a 2-tailed p-value and a Pearson correlation coefficient.

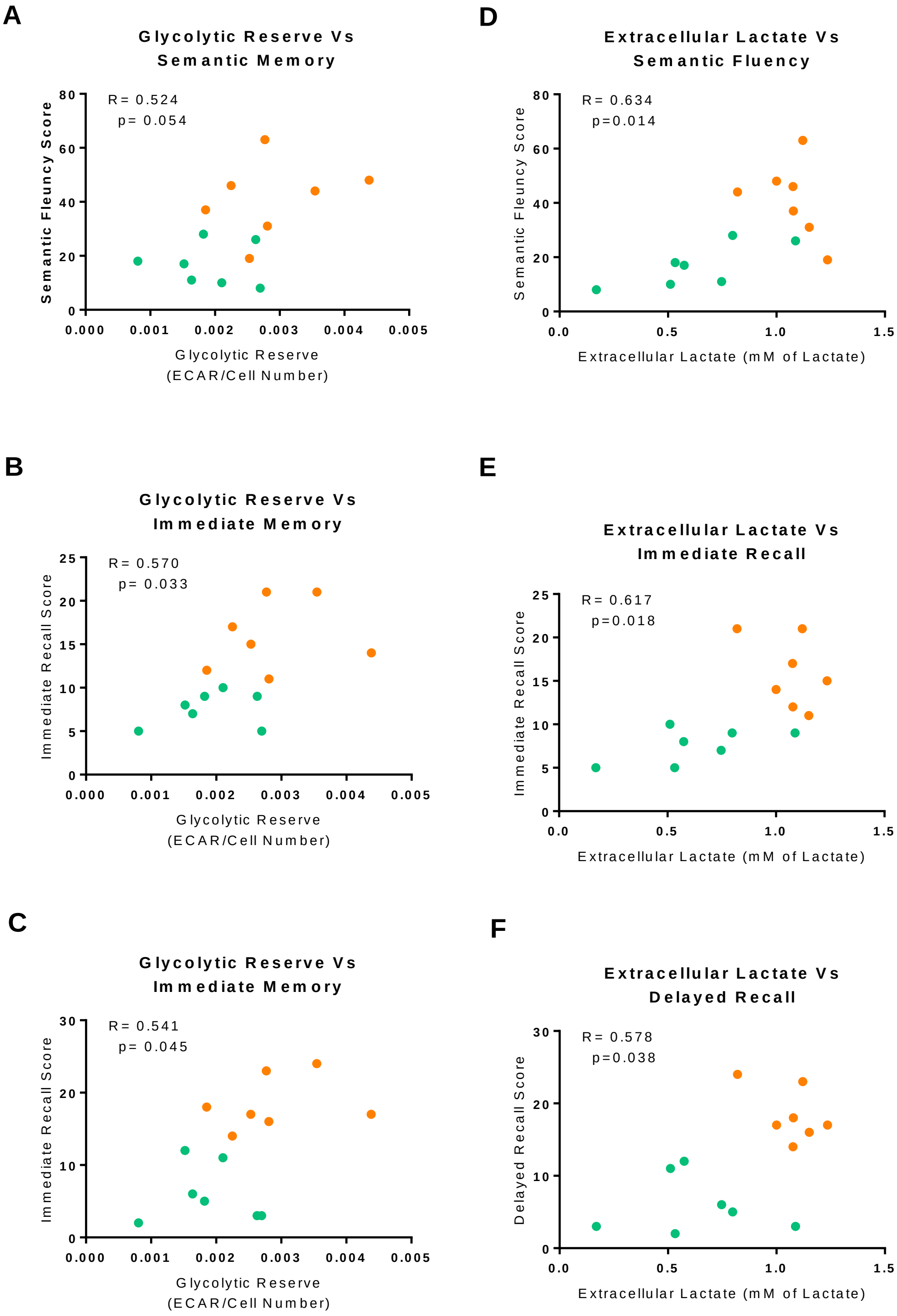


**Supplementary Figure 3 |** ***Glycolytic reserve and extracellular lactate correlations with Neuropsychological tests*** This figure highlights the glycolytic reserve correlations with semantic memory (Figure 3A) immediate recall (Figure 3B) and delayed recall (Figure 3C).

Astrocyte extracellular lactate correlations with semantic memory (Figure 3D) immediate recall (Figure 3E) and delayed recall (Figure 3F) are also displayed. Green points represent sporadic AD patients and orange points represent control subjects. In each correlation 7 controls and 5 sAD cell lines are plotted. The correlations are performed using a 2-tailed p-value and a Pearson correlation coefficient.

| Neuropsychological Tests | Astrocyte Metabolic Marker | R-Value | P-value |
|---|---|---|---|
| Immediate Recall | | | |
| | Mitochondrial Spare Respiratory Capacity | 0.586 | 0.075 |
| Immediate Recall | | | |
| | Glycolytic Reserve | 0.824 | **0.003** |
| Delayed Recall | | | |
| | Glycolytic Reserve | 0.658 | **0.039** |
| Immediate Recall | | | |
| | Extracellular Lactate Level | 0.552 | 0.123 |
| Delayed Recall | | | |
| | Extracellular Lactate Level | 0.368 | 0.330 |
| Semantic Fluency | | | |
| | Extracellular Lactate Level | 0.865 | **0.003** |

**Supplementary Table 3 |** ***Astrocyte neuropsychological metabolic correlations*** This table displays the correlations between different astrocyte metabolic parameters and neuropsychological testing after controlling for participant age, brain reserve and length of education. For this analysis the controls and sAD astrocytes were considered as one group. Numbers highlighted in bold are statistically significant correlations.

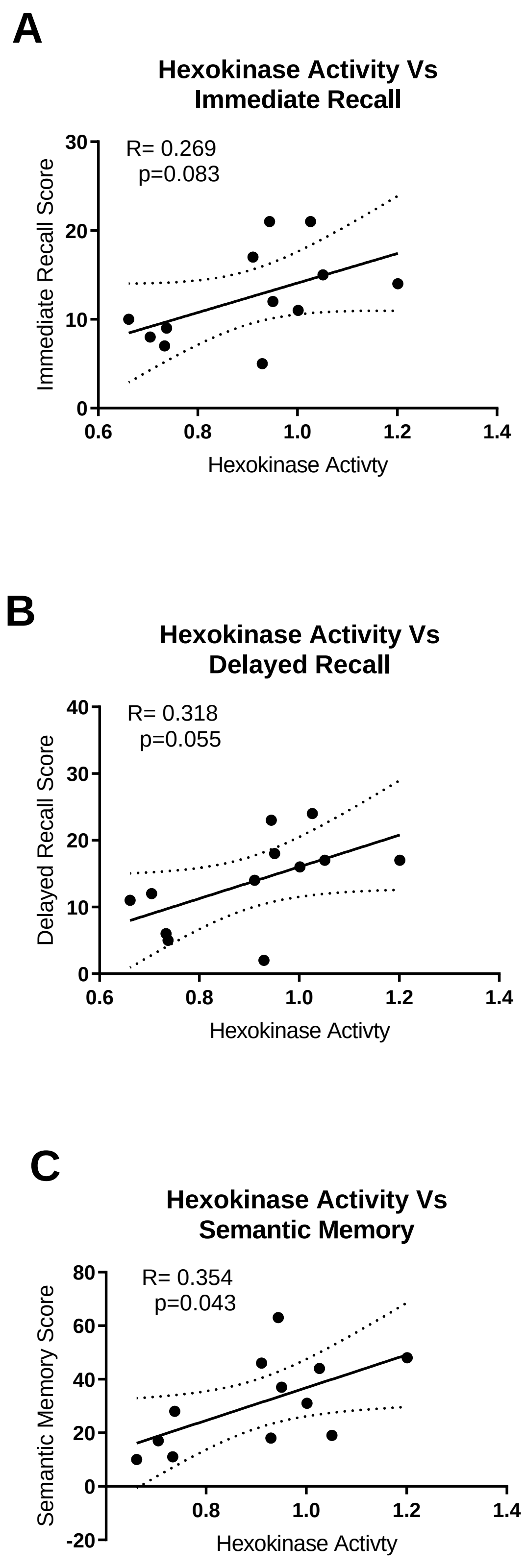
A
Hexokinase Activity Vs
Immediate Recall
R= 0.269
p=0.083
Immediate Recall Score
30
20
10
0
0.6
0.8
1.0
1.2
1.4
Hexokinase Activty
B
Hexokinase Activity Vs
Delayed Recall
R= 0.318
p=0.055
Delayed Recall Score
40
30
20
10
0
0.6
0.8
1.0
1.2
1.4
Hexokinase Activty
C
Hexokinase Activity Vs
Semantic Memory
R= 0.354
p=0.043
Semantic Memory Score
80
60
40
20
0
-20
0.8
1.0
1.2
1.4
Hexokinase Activty

**Supplementary Figure 4 |** ***Hexokinase 1 correlations with neuropsychological tests.*** In each correlation 7 controls and 5 sAD cell lines are plotted. The correlations are performed using a 2-tailed p-value and a Pearson correlation coefficient.
